# The Genetics of Aerotolerant Growth in a Naturally Reduced Genome Alphaproteobacterium

**DOI:** 10.1101/2023.06.12.544693

**Authors:** Amy L. Enright, Amy B. Banta, Ryan D. Ward, Julio Rivera Vazquez, Magdalena M. Felczak, Michael B. Wolfe, Michaela A. TerAvest, Daniel Amador-Noguez, Jason M. Peters

## Abstract

Reduced genome bacteria are genetically simplified systems that facilitate biological study and industrial use. The free-living Alphaproteobacterium, *Zymomonas mobilis*, has a naturally reduced genome containing fewer than 2000 protein coding genes. Despite its small genome, Z. mobilis thrives in diverse conditions including the presence or absence of atmospheric oxygen. However, insufficient characterization of essential and conditionally essential genes has limited broader adoption of *Z. mobilis* as a model Alphaproteobacterium. Here, we use genome-scale CRISPRi-seq to systematically identify and characterize *Z. mobilis* genes that are conditionally essential for aerotolerant or anaerobic growth, or are generally essential across both conditions. Comparative genomics revealed that the essentiality of most “generally essential” genes was shared between *Z. mobilis* and other Alphaproteobacteria, validating *Z. mobilis* as reduced genome model. Among conditionally essential genes, we found that the DNA repair gene, recJ, was critical only for aerobic growth but reduced the mutation rate under both conditions. Further, we show that genes encoding the F_1_F_O_ ATP synthase and Rnf respiratory complex are required for anaerobic growth of *Z. mobilis*. Combining CRISPRi partial knockdowns with metabolomics and membrane potential measurements, we determined that the ATP synthase generates membrane potential that is consumed by Rnf to power downstream processes. Rnf knockdown strains accumulated isoprenoid biosynthesis intermediates, suggesting a key role for Rnf in powering essential biosynthetic reactions. Our work establishes *Z. mobilis* as a streamlined model for alphaproteobacterial genetics, has broad implications in bacterial energy coupling, and informs *Z. mobilis* genome manipulation for optimized production of valuable isoprenoid-based bioproducts.

**Importance:** The inherent complexity of biological systems is a major barrier to our understanding of cellular physiology. Bacteria with markedly fewer genes than their close relatives, or reduced genome bacteria, are promising biological models with less complexity. Reduced genome bacteria can also have superior properties for industrial use, provided the reduction does not overly restrict strain robustness. Naturally reduced genome bacteria, such as the Alphaproteobacterium, *Zymomonas mobilis*, have fewer genes but remain environmentally robust. In this study, we show that *Z. mobilis* is a simplified genetic model for Alphaproteobacteria, a class with important impacts on the environment, human health, and industry. We also identify genes that are only required in the absence of atmospheric oxygen, uncovering players that maintain and utilize the cellular energy state. Our findings have broad implications for the genetics of Alphaproteobacteria and industrial use of *Z. mobilis* to create biofuels and bioproducts.

## Introduction

Reduced genome bacteria are powerful models for dissecting biological function and streamlined platforms for industrial production. These bacteria have less of the genetic redundancy that often obscures the functions of genes and pathways, simplifying modeling of cellular processes (1, 2). Bacteria with experimentally reduced genome sizes have shown improved properties for industrial applications such as increased genomic stability (3, 4), faster growth rates (5), improved transformation efficiency (6), and ease of genetic manipulation (7), all of which facilitate introduction and maintenance of engineered pathways. However, experimental genome reduction can also result in substantial growth defects (8, 9) and loss of robustness to environmental conditions (10) if the physiology of the bacterium is not well understood and taken into consideration. An appealing alternative approach would be to utilize bacteria with naturally reduced genomes compared to related species. Such bacteria would effectively be “pre-evolved” with the benefits of a reduced genome, but with the robustness of environmental strains.

*Zymomonas mobilis* is an emerging model for bacteria with naturally reduced genomes and has excellent properties as an industrial platform. *Z. mobilis* is a member of the highly-studied class, Alphaproteobacteria, but has at least 1000 fewer genes than closely related species (a total of 1915 protein coding genes (11, 12)). This natural genome reduction has created a streamlined metabolism that efficiently ferments sugars to ethanol (13, 14), aiding metabolic modeling efforts (15) and highlighting the promise of *Z. mobilis* as a biofuel producer. Despite its reduced genome, *Z. mobilis* is free-living and grows quickly to high densities in standard rich medium (*e.g.*, buffered yeast extract and glucose (16)). In contrast, other reduced genome models are often endosymbionts with fastidious growth requirements (17). *Z. mobilis* growth is robust to diverse environmental conditions including the presence or absence of atmospheric oxygen (aerotolerant anaerobe), high concentrations of ethanol (up to 16% (v/v)) (18), and some but not all inhibitors found in plant-derived biofuel fermentation substrates (19). Its safety profile (Generally Regarded As Safe (20)), ease of manipulation in aerobic settings (21), and excellent anaerobic fermentation properties (14) indicate that *Z. mobilis* is both an outstanding model for basic biology of Alphaproteobacteria and a rising industrial workhorse.

Genes required for robust growth of *Z. mobilis* across conditions are understudied, hindering both its use as a reduced genome model and rational engineering efforts to optimize biofuel production. In these contexts, generally and conditionally essential genes are of particular interest because they must be maintained by reduced genomes and are linked to core cellular processes such as carbon metabolism that impact biofuel production (15). Moreover, reduced genomes are thought to harbor a larger fraction of essential genes than larger genomes (17), underscoring the importance of such genes in bacteria with less genetic redundancy. Although transposon (Tn) mutagenesis is often used at the genome-scale to identify essential genes (22, 23), previous attempts to apply this approach to *Z. mobilis* were unsuccessful (24), possibly due to the polyploid nature of the *Z. mobilis* chromosome (25–27). Regardless, Tn or other gene disruption approaches alone cannot be used to phenotype genes in the condition where they are essential, since this results in cell death.

With the goal of defining and characterizing essential genes, we previously developed a CRISPRi (<u>c</u>lustered regularly interspaced <u>p</u>alindromic repeats interference) gene knockdown system for *Z. mobilis* (28). CRISPRi targets genes for knockdown using a single guide (sg)RNA which directs a catalytically dead (d)Cas9 nuclease to a complementary gene target where the sgRNA-dCas9 complex binds and blocks transcription (29). Our *Z. mobilis* CRISPRi system has several important advantages: it is IPTG inducible, titratable with subsaturating inducer or using mismatched sgRNAs (30), stably integrated into the chromosome without the need for selection, and knocks down multi-copy genes simultaneously—features which enable interrogation of both non-essential and essential genes in this plausibly polyploid bacterium.

Here, we combine comparative and functional genomics approaches to establish *Z. mobilis* as a genetic model for Alphaproteobacteria. We use genome-scale CRISPRi to identify genes that are generally or conditionally essential depending on the presence of environmental oxygen. We find that generally essential *Z. mobilis* genes represent a core set of genes that are highly conserved across Alphaproteobacteria. Our sets of *Z. mobilis* aerobic and anaerobic essential genes contain several surprising players, revealing an oxygen-dependent requirement for the RecJ DNA repair protein and critical roles for the ATP synthase and Rnf complex in maintence and utilization of the proton motive force (PMF) during anaerobic growth. Our studies provide a genetic window into how a naturally reduced genome can be both streamlined and robust.

## Results

### *Z. mobilis* is an emerging model for small-genome, free-living microorganisms

A defining feature of *Z. mobilis* is its naturally small genome of only 1915 protein coding genes, which are encoded across one ∼2.06 Mb chromosome and four endogenous plasmids (0.03-0.04 Mb each) (12). In contrast, the vast majority of free-living Alphaproteobacteria carry many additional genes (Fig. 1A), including some of the closest *Z. mobilis* relatives, such as *Sphingomonas wittichii* and *Rhodobacter sphaeroides* (∼5300 and ∼4200 genes, respectively) (Fig. 1B) (31, 32). Endosymbiont Alphaproteobacteria (*e.g.*, *Rickettsia* and *Wolbachia*) (Fig. 1A) have even fewer genes than *Z. mobilis* but rely on their host to complement various missing gene functions (33, 34).

**Figure 1.**
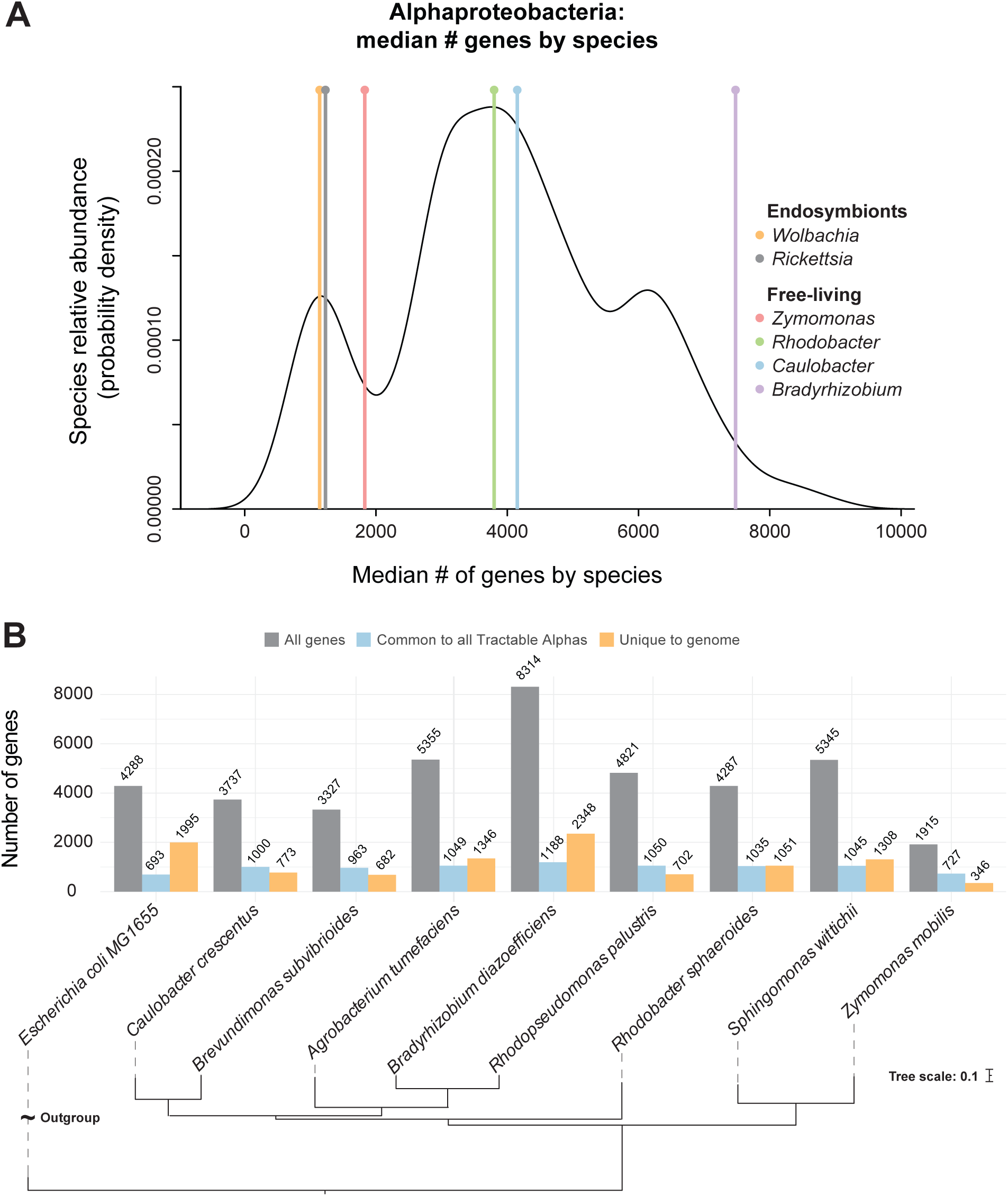
*Zymomonas mobilis* as a small-genome model for Alphaproteobacteria. (A) Density plot displaying median number of genes by species of Alphaproteobacteria. Vertical lines mark median number of genes per genus noted. (B) Gray lefthand bars; number of genes per organism. Blue middle bars; number of genes per organism with homologs in all seven Tractable Alphas (*C. crescentus, B. subvibrioides, A. tumefaciens, B. diazoefficiens, R. palustris, R. sphaeroides, S. wittichii*). Note these numbers encompass homologous genes within each organism and thus vary slightly across the Tractable Alphas. Orange righthand bars; number of genes per organism with no homologs in any of the seven Tractable Alphas. Phylogenetic tree generated with OrthoFinder using annotated protein-coding genes.

To evaluate *Z. mobilis* as a model Alphaproteobacterium, we compared the *Z. mobilis* strain ZM4 genome against seven genetically tractable Alphaproteobacteria (“Tractable Alphas”) chosen based on availability of both a genome sequence and an essential gene list curated from Tn screens (35–40). Given the low genetic diversity within the genus *Zymomonas* (16), ZM4 serves as a strong representation of the genus overall. We found that of the 890 protein-coding genes that are conserved across all seven Tractable Alphas, *Z. mobilis* has orthologs for 701 (∼79%) (Fig. S1, Table S8). As expected, this group of genes was enriched for core cellular processes such as translation (FDR = 2.9e^-7^) and central metabolic processes (FDR = 9.1e^-6^). *Z. mobilis* also carries relatively few unique genes that do not share orthologs in any of the Tractable Alphas (346 unique genes, fewer than any of the Tractable Alphas) (Fig. 1B); these genes largely encode horizontally-transferred elements (e.g., CRISPR-associated immune systems) and hypothetical proteins (Table S8). Genes carried by other Alphaproteobacteria but not by *Z. mobilis* are largely related to oxidative phosphorylation (*e.g.*, type I NADH dehydrogenase, cytochrome c oxidase) and some TCA cycle components that are known to be absent in *Z. mobilis* (Table S8) (41–45).

### A whole-genome CRISPRi library for *Z. mobilis*

To interrogate the genetic requirements of this small-genome model species, we constructed a whole-genome knockdown library using our previously developed *Z. mobilis* CRISPRi system (46, 47), a task made further feasible by its minimalistic genome. For nearly all *Z. mobilis* genes (∼98%, 1895/1933), we successfully designed and cloned four corresponding sgRNAs intended to provide strong knockdown (*i.e.*, the sgRNA sequence is fully complementary to the gene of interest). For the remaining 38 genes, the library contains 1-3 sgRNAs per gene. When assessing gene-level fitness, we considered the behavior of the median sgRNA, a strategy which aims to neutralize any biases incurred from ineffective, off-target, or “bad seed” sgRNAs (48). The library also contains 1000 non-targeting sgRNAs that act as internal controls.

CRISPRi is inherently polar, where knockdown of an upstream gene within a transcription unit (TU) causes concurrent knockdown of the downstream gene(s) (29, 49). Conversely, “reverse polarity” is a poorly characterized CRISPRi phenomenon in which knockdown of a downstream gene causes decreased expression of the upstream gene(s) (50, 51). Previous studies have observed reverse polarity in *Bacillus subtilis* (49) and to a lesser extent in *Escherichia coli* (48). In contrast, reverse polarity was largely undetected in *Mycobacterium tuberculosis* (52). To investigate if reverse polarity occurs in *Z. mobilis*, we designed sgRNAs targeted along a reporter TU encoding two fluorescent proteins. Our findings revealed that reverse polarity is indeed present in *Z. mobilis*, as knockdown of the downstream sfGFP-encoding gene resulted in decreased expression of the upstream mScarlet reporter, with the strongest reverse-polar effects observed when targeting nearest to the 5’ end of the downstream gene (5.4-fold mScarlet knockdown when targeting the 5’ end of sfGFP and 1.8-fold when targeting 3’ end) (Fig. S2). Similar results were observed in *E. coli* (3.1-fold mScarlet knockdown when targeting the 5’ end of sfGFP and no detectable mScarlet knockdown when targeting 3’ end of sfGFP) (Fig. S2). Together, these results underscore the known importance of interpreting CRISPRi results at the level of TUs. To account for this, we note predicted TUs (Table S7) when reporting gene phenotypes.

### Comprehensive gene function interrogation across oxygen diverse environments

To understand how *Z. mobilis* employs its small genome to flourish in oxygen diverse environments, we utilized the whole-genome CRISPRi library to identify genes that are conditionally essential for aerobic or anaerobic growth and genes that are generally essential regardless of condition. Library cultures were grown with saturating concentrations of IPTG (1 mM) to induce CRISPRi and were maintained in exponential phase for ∼10 doublings at which point samples were taken to measure sgRNA spacer abundance by Illumina sequencing (Fig. 2A).

**Figure 2.**
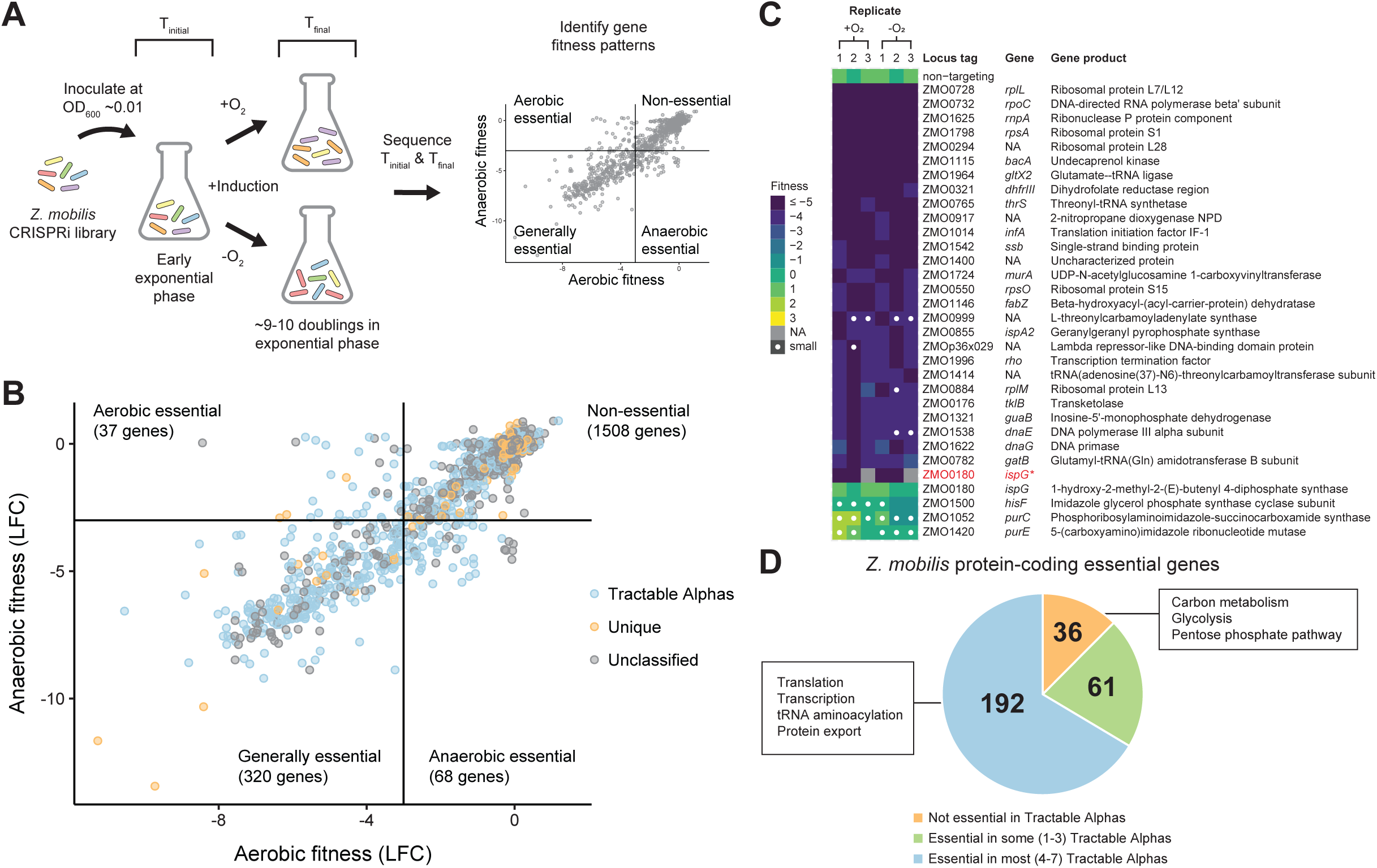
Comprehensive interrogation of gene function and essentiality in *Zymomonas mobilis* using CRISPRi. (A) Overview of whole-genome CRISPRi screen to assay *Z. mobilis* gene function during aerobic versus anaerobic growth. (B) *Z. mobilis* genes classified as conditionally essential for aerobic versus anaerobic growth, generally essential, or non-essential. LFC, median log2 fold-change for sgRNAs targeting a particular gene for knockdown. Blue, *Z. mobilis* genes with homologs in all seven Tractable Alphas (*C. crescentus, B. subvibrioides, A. tumefaciens, B. diazoefficiens, R. palustris, R. sphaeroides, S. wittichii*); orange, genes unique to *Z. mobilis* genome, defined by sharing no homologs in any of the seven Tractable Alphas; gray, remaining unclassified *Z. mobilis* genes. (C) Validation of generally essential gene phenotypes using a new validation sgRNA that was not included in the CRISPRi library. Heat map displays fitness scores for knockdown mutants assayed individually on spot plates. White dots denote abnormally small colonies (see Methods). Asterisk marks repeated validation of *ispG* using an sgRNA from our CRISPRi library with an LFC close to the gene median, rather than a new validation sgRNA. Two replicates were assayed for the repeat validation of *ispG*. (D) Of the 289 protein-coding *Z. mobilis* essential genes, 192 (∼66%) share essential homologs in most (4–7) of the Tractable Alphas, 61 (∼21%) share essential homologs in some (1–3) of the Tractable Alphas, and 36 (∼12%) share no essential homologs with any of the Tractable Alphas.

Likely due to its small genome size, *Z. mobilis* harbors a larger fraction of essential genes than other model organisms. We identified 320 genes that were essential across both conditions, conservatively defined by log_2_ fold change (LFC) ≤ −3 and significance ≤ 0.05 (Fig. 2B, Table S9). A similar approach (i.e., LFC < −2) recovered 79% of known *E. coli* essential genes from a genome-scale CRISPRi screen (53), suggesting that our cutoffs largely guard against false positives. Of the 320 generally essential *Z. mobilis* genes, 289 are protein-coding; remarkably, these genes represent ∼15% of the *Z. mobilis* genome. In comparison, the *E. coli* and *B. subtilis* genomes are made up of only ∼7% (307/4131) and ∼6% (257/4245) protein-coding essential genes, respectively (54–56). Thus, *Z. mobilis* models the unique features of small-genome, free-living species in a way that other model bacteria cannot.

To verify these phenotypes outside of the pooled library context, we chose 31 generally essential genes to assay individually using a new validation sgRNA that was not included in the library. Each knockdown strain was serially diluted and spotted onto plates containing IPTG inducer alongside a non-targeting control then analyzed following growth for plating and colony size defects (see Methods). Initially, we successfully recapitulated phenotypes for 27 of the 31 essential gene knockdowns tested (Fig. 2C, Fig. S3). Non-recapitulated phenotypes may result from false positives, specificity to the pooled context, differences in essentiality phenotype during growth on a plate versus in liquid culture, or ineffective validation sgRNAs. For example, we were at first unable to validate the generally essential gene *ispG* (ZMO0180) using the new validation sgRNA. However, when we instead used an sgRNA from our library with an LFC close to the gene median, we successfully validated the *ispG* phenotype outside of the pooled context (Fig. 2C, Fig. S3), bringing the total recapitulated phenotypes to 28 of 31 (∼90%).

Comparison of the gene essentiality profile of *Z. mobilis* to the seven Tractable Alphas further solidified its value as a model, small-genome Alphaproteobacterium. We found that ∼88% of the 289 generally essential *Z. mobilis* genes have homologs that are also essential in at least one other Alphaproteobacterium, with ∼66% sharing essential homologs in most (≥4) Tractable Alphas (Fig. 2D, Table S8), thus demonstrating substantial agreement between core essential genes of *Z. mobilis* and other Alphaproteobacteria. As expected, essential genes conserved across most Tractable Alphas included fundamental cellular processes such as transcription, translation and protein export. Genes uniquely essential in *Z. mobilis* were functionally enriched for carbon metabolism (*e.g.*, glycolysis and the pentose phosphate pathway), reflecting a low redundancy, “catabolic highway” (57) ideal for industrial engineering. Other unique essentials included plasmid-borne, predicted DNA-binding proteins. Given that the native *Z. mobilis* plasmids can be cured without loss of viability (58), we speculate that these proteins are negative regulators of toxic genes that contribute to plasmid maintenance.

### Oxygen-dependent roles for oxygen-resistant enzymes, DNA repair, and ferredoxin/flavodoxin-based redox systems

Tn libraries are often constructed aerobically, obscuring the distinction between genes that are conditionally essential for aerobic growth versus those that are generally essential. In contrast, CRISPRi libraries can be generated aerobically without perturbing genes during the construction process, enabling discovery of genes that are conditionally essential for aerobic growth. Following the same quantitative approach as above, we next identified 37 conditionally essential genes for aerobic growth (LFC ≤ −3 and significance ≤ 0.05 for aerobic growth but not for anaerobic growth) (Fig. 2B, Fig. 3A, Table S9). In multiple cases, *Z. mobilis* has redundant enzymes performing similar functions, except one is oxygen-tolerant and the other is not. For example, *nrdA* (TU: ZMO1039) and *nrdF* (TU: ZMO0443), which encode class I ribonucleotide reductases (RNRs) that synthesize deoxyribonucleotides for DNA replication and repair, are conditionally essential for aerobic growth, whereas class III RNRs are oxygen-sensitive (59). Similarly, genes encoding the pyruvate dehydrogenase complex, which converts pyruvate to acetyl-CoA aerobically or anaerobically (60), including *pdhAB* (TU: ZMO1605-1606; *pdhA, pdhB*), *pdh*C and *lpd* (TU: ZMO0510-ZMO0513; *pdhC*, ZMO0511, *lpd*, ZMO0513) are conditionally essential for aerobic growth because the other enzyme responsible for this function (pyruvate formate lyase) is oxygen-sensitive (61). The presence of redundant aerobic essentials in an otherwise streamlined genome points to a key role for aerotolerant growth in the natural environment of *Z. mobilis*.

**Figure 3.**
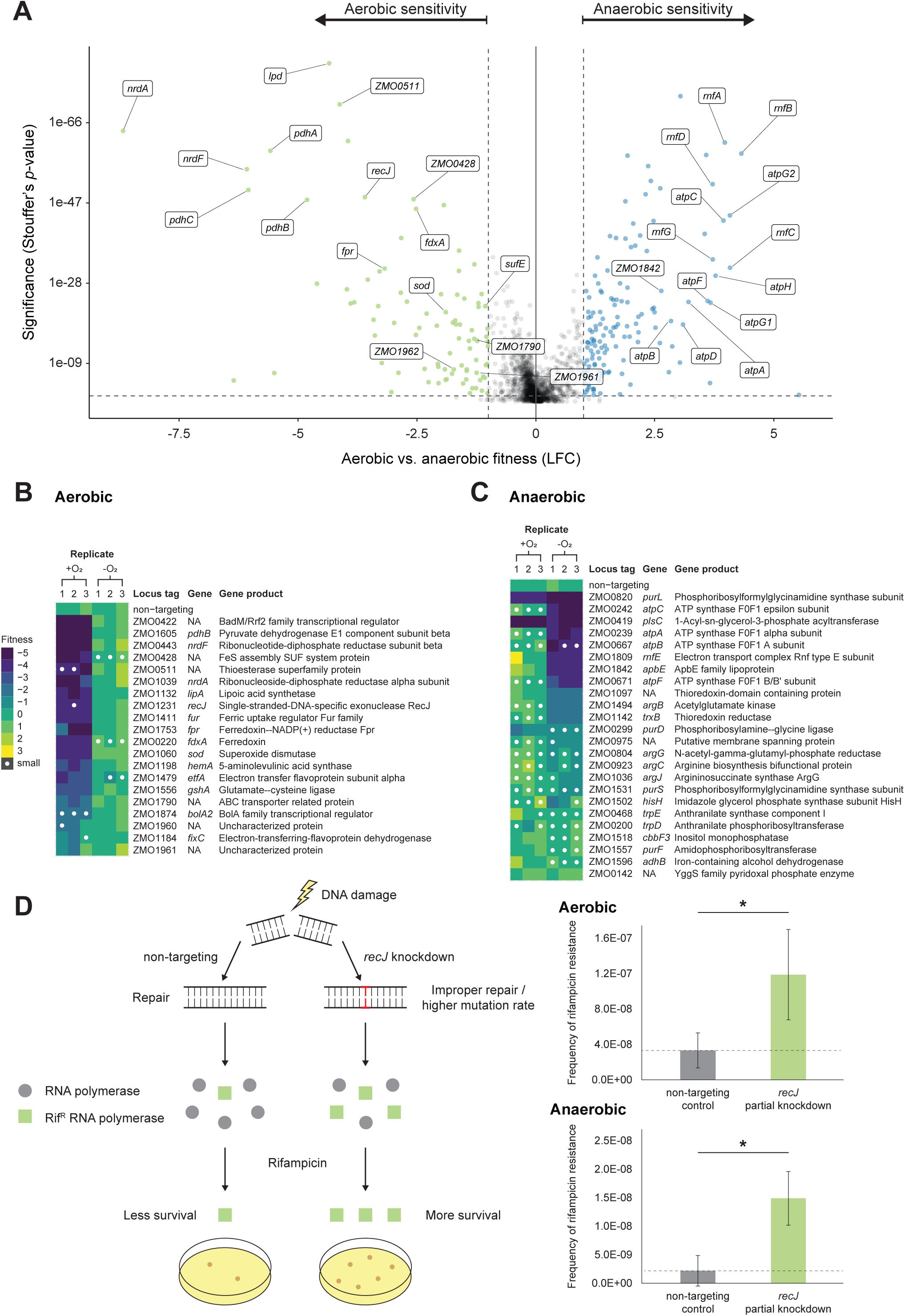
Whole-genome CRISPRi screen reveals condition-specific gene phenotypes for aerobic versus anaerobic growth of *Z. mobilis*. (A) Genes with aerobic or anaerobic sensitivity, defined by a fitness score (log2 fold-change, LFC) ≥ |1| and significance (Stouffer’s *p*-value) < 0.05. Dashed lines depict these fitness and significance cutoffs. Green dots, genes with aerobic sensitivity. Blue dots, genes with anaerobic sensitivity. (B-C) Validation of (B) aerobic-specific sensitivity and (C) anaerobic-specific sensitivity gene phenotypes using a new validation sgRNA that was not included in the CRISPRi library. Heat maps display fitness scores for knockdown mutants assayed individually on spot plates. White dots denote abnormally small colonies (see Methods). (D) Left, schematic depicting rifampicin assay as a proxy for mutation rate. In this assay, improper DNA repair elevates the mutation rate and yields increased abundance of surviving rifampicin-resistant (RifR) mutants. Right, frequency of rifampicin resistance for *recJ* partial knockdown versus non-targeting CRISPRi control. Top right, aerobic. Bottom right, anaerobic. Asterisk denotes significance (*p* < 0.01) by two-tailed Student’s T-test. Data represent three replicates.

Consistent with previous work that identified *Z. mobilis* genes with increased expression following a shift from anaerobic to oxygen-replete conditions (21), knockdown of known oxidative stress genes such as *sod* (superoxide dismutase; TU: ZMO1060) and iron-sulfur cluster assembly (*suf*) genes including *sufE* (TU: ZMO1067-1068; *sufE*, ZMO1068), ZMO0422 (homolog of transcriptional regulator IscR), and ZMO0428 (TU: ZMO0422-0429; ZMO0422, *sufB, sufC, sufD, sufS*, ZMO0428, ZMO0429) also showed oxygen-specific fitness defects (Fig. 3A, Fig. 3B, Table S9).

Interestingly, the DNA repair gene *recJ* (TU: ZMO1231) was also found to be conditionally essential for aerobic growth, and this phenotype was validated outside the pooled context (Fig. 3A, Fig. 3B). Although non-essential in *E. coli* (62), previous work has shown that *recJ* is essential across the Alphaproteobacteria (63) and in *Deinococcus radiodurans* (64, 65) for aerobic growth. However, these studies did not (or could not, in the case of strict aerobes) compare the *recJ* phenotype during aerobic versus anaerobic growth. *recJ* knockdowns in *Z. mobilis* showed no apparent growth phenotype under anaerobic conditions. To understand the role of RecJ in Alphaproteobacterial physiology, we examined the mutation frequency in *recJ* knockdown strains. We found that RecJ suppresses the mutation frequency in both aerobic and anaerobic conditions. Because full knockdown of *recJ* is lethal aerobically, we used a subsaturating concentration of inducer to create partial knockdowns. To calculate mutation frequency, we measured the number of rifampicin-resistant colony forming units in non-targeting and *recJ* partial knockdown strains. Knockdown of *recJ* increased the mutation frequency in both aerobic (3.6-fold) and anaerobic conditions (6.8-fold; Fig. 3D); thus, we establish a role for RecJ in mutation suppression in *Z. mobilis* that may be applicable to other Alphaproteobacteria. However, the mechanistic rationale for *recJ* essentiality during aerobic growth remains elusive and requires further studies. We were additionally able to associate an oxygen-specific phenotype with unannotated or poorly annotated genes including the TU encoding ZMO1961-ZMO1962 (uncharacterized protein and putative histidine kinase, respectively), and ZMO1790 (TU: ZMO1790; predicted heme-related transporter (66)) (Fig. 3A, Fig. 3B, Table S9). Finally, we identified ferredoxin/flavodoxin (Fd/Fld)-based redox systems specifically required for aerobic growth including *fdxA* (TU: ZMO1753) and *fpr* (Fd/Fld reductase; TU: ZMO0220). As we will highlight in the next section, such redox functions play a similarly indispensable role in anaerobic growth, though the relevant genetic players may differ by condition.

### Rnf respiratory complex and F_1_F_O_ ATP synthase are essential for anaerobic growth

We next examined the 68 genes that were conditionally essential for growth in the absence of oxygen (LFC ≤ −3 and significance ≤ 0.05 for anaerobic growth but not for aerobic growth) (Table S9). Strikingly, TUs encoding either the Rnf complex (TU: ZMO1808-1814; *rnfH, rnfE, rnfG, rnfD, rnfC, rnfB, rnfA*) or the F_1_F_O_ ATP synthase (TU-1: ZMO0238-0241; *atpH, atpA, atpG1, atpD*). TU-2: ZMO0242; *atpC*. TU-3: ZMO0669-0671; *atpG2, atpF*) were conditionally essential for anaerobic growth. Of note, while *rnf* knockdown was inconsequential for growth in the presence of oxygen (median aerobic LFC for *rnf* genes = −0.2), *atp* knockdown resulted in a mild aerobic fitness defect (median aerobic LFC for *atp* genes = −1.2) (Table S9). The F_1_F_O_ ATP synthase reversibly couples H^+^ transport with the synthesis or hydrolysis of ATP. Rnf, originally named for its role in *<u>R</u>hodobacter* <u>n</u>itrogen fixation, is similarly reversible and couples ion transport with electron transfer between ferredoxin and pyridine nucleotides (67). A previous Tn study by Deutschbauer, *et al.* also observed that *rnf* genes are required for anaerobic growth of *Z. mobilis*; however, the mechanism underlying this phenotype was not explored further (66). The same group proposed ZMO1842 to be *rnfF* (TU: ZMO1842-1842; possibly *rnfF*, *rseC*) based on cofitness with other *rnf* genes. This gene was also conditionally essential for anaerobic growth in our screen (Table S9), but it presently is not annotated as *rnfF* and thus was not included in our subsequent *rnf* analyses.

We sought to understand the potential mechanism driving the *rnf* and *atp* anaerobic growth phenotypes. First, we tested whether the complexes were functioning in an ion gradient-dependent manner in *Z. mobilis*. Dependency on the proton gradient was tested using carbonyl cyanide 3-chlorophenylhydrazone (CCCP), a protonophore that dissipates PMF. We reasoned that partial knockdown of *rnf* and *atp* genes could act as a sensitized genetic background against which to test CCCP challenge. Given that the *atp* genes are encoded across three operons in *Z. mobilis*, we targeted one gene per *atp* operon for knockdown (*atpA, atpC, atpF*). For *rnf* knockdown, *rnfE* was targeted. Partial knockdown was induced with a subsaturating concentration of IPTG inducer. We found that, under anaerobic conditions, *rnf* and *atp* knockdowns had increased sensitivity to a normally sub-lethal concentration of CCCP, demonstrating their interdependence on PMF (Fig. 4A). Consistent with our screen results, when the assay was repeated aerobically, the *rnf* knockdown mutant behaved like the non-targeting control while *atp* knockdowns retained some sensitivity to CCCP, though to a lesser extent than was seen in the absence of oxygen (Fig. 4A).

**Figure 4.**
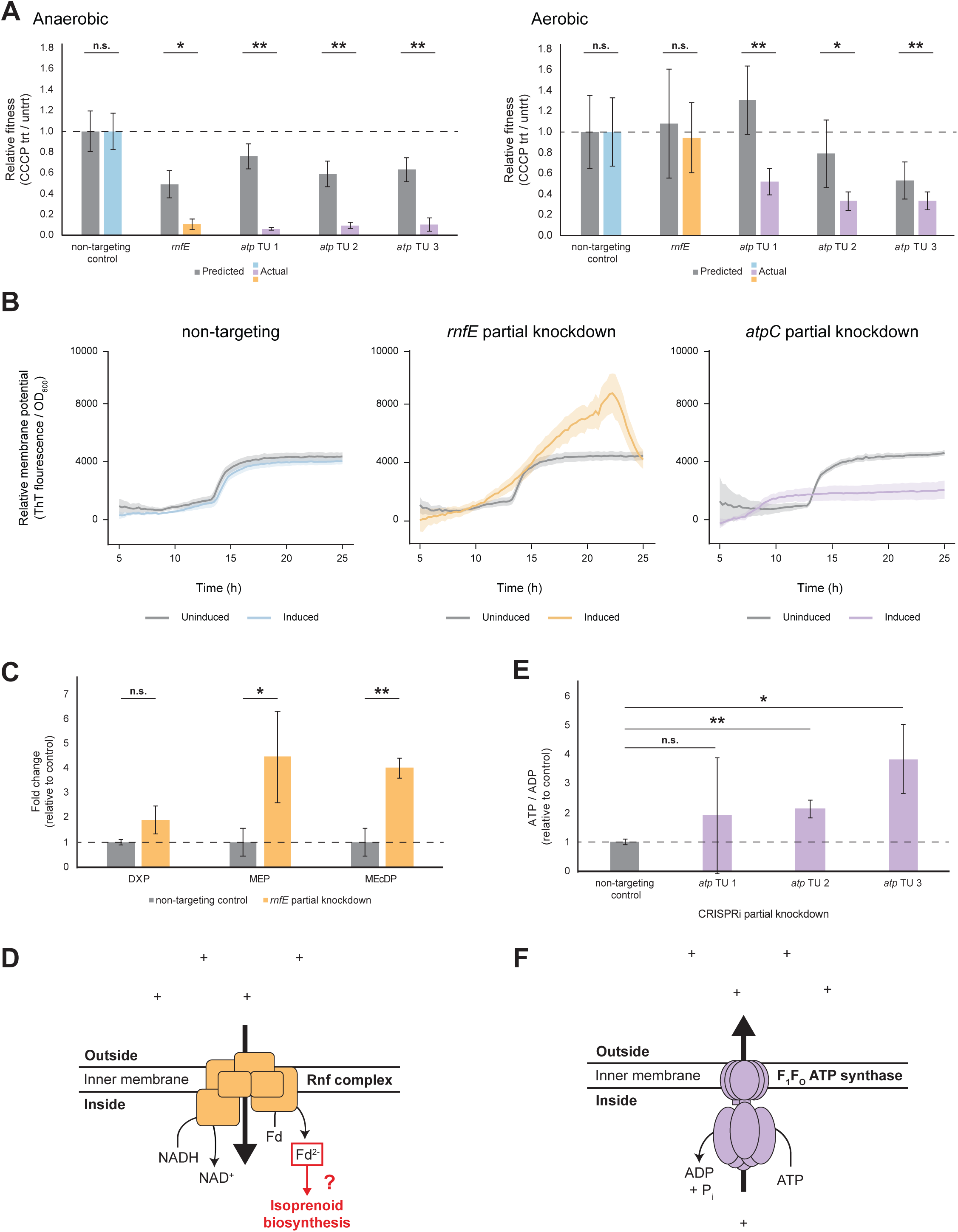
Functional analysis of the *Z. mobilis* F_1_F_O_ ATP synthase and Rnf complex during anaerobic growth. n.s., not significant; *, *p* < 0.05; **, *p* < 0.01 by two-tailed Student’s T-test. TU, transcription unit. *atp* TU 1 knockdown targets *atpA*; *atp* TU 2 knockdown targets *atpC*; *atp* TU 3 knockdown targets *atpF*. (A) Fitness of *rnf* or *atp* gene knockdowns during disruption of the electrochemical gradient by sublethal challenge with the protonophore CCCP. Fitness values are relative to the non-targeting control (horizontal dashed line). Gray lefthand bars represent predicted fitness by the multiplicative model (fitness in CCCP × fitness of partial knockdown). Colored righthand bars show measured fitness of non-targeting control (blue), *atp* knockdowns (purple), or *rnf* knockdown (orange). Left, anaerobic. Right, aerobic. Data represent three experiments with two biological replicates each. (B) Thioflavin T (ThT) assay to measure relative membrane potential of (left) non-targeting CRISPRi control, (middle) *rnfE* partial knockdown, or (right) *atpC* partial knockdown. Gray lines, uninduced; colored lines, induced partial knockdown. Shaded ribbons represent one standard deviation from the mean. Data represent six replicates. (C) Mass spectroscopy metabolomics measurement of MEP pathway intermediates for (lefthand gray bars and dashed line) non-targeting CRISPRi control and (righthand orange bars) *rnfE* partial knockdown. DXP, 1-deoxy-D-xylulose 5-phosphate; MEP, 2-C-methyl-D-erythritol 4-phosphate; MEcDP, 2-C-methyl-D-erythritol-2,4-cyclodiphosphate. Data represent three replicates. (D) Speculative model for Rnf function during Z. mobilis anaerobic growth: Rnf utilizes the ion motive force to reduce ferredoxin, and reduced ferredoxin (Fd^2-^) powers important biological processes such as isoprenoid biosynthesis via the MEP pathway. (E) Mass spectroscopy metabolomics measurement of ATP/ADP ratio for (gray bars and dashed line) non-targeting CRISPRi control and (purple bars) *atp* partial knockdowns. Data represent three replicates. (F) Model for F_1_F_O_ ATP synthase function during *Z. mobilis* anaerobic growth: ATP is hydrolyzed to pump ions across the inner membrane.

Given that the reactions carried out by both complexes are reversible, multiple models could explain the synergy of the proton gradient with F_1_F_O_ ATP synthase and Rnf. In each case, interdependence with the ion gradient could indicate either that the complex functions to establish the gradient (a producer), or that it depends on the gradient to carry out its essential function (a consumer). Thus, we next investigated the directionality of both complexes during anaerobic growth.

Two lines of evidence support the hypothesis that Rnf operates as a membrane potential consumer during anaerobic growth. First, a Thioflavin T (ThT) membrane potential assay, wherein increased ThT fluorescence indicates stronger membrane potential (68, 69), revealed that partial knockdown of *rnfE* yielded stronger membrane potential than an uninduced control (Fig. 4B). Second, mass spectrometry metabolomics analysis of the *rnfE* partial knockdown uncovered an accumulation of intermediate metabolites in the methylerythritol 4-phosphate (MEP) isoprenoid biosynthesis pathway. Iron-sulfur cluster enzymes involved in the MEP pathway require reduced ferredoxin/flavodoxin (Fd_red_/Fld_red_) (21). We observed ≥4-fold accumulation of two intermediates, MEP and MEcDP (Fig. 4C). Thus, we infer that *rnfE* knockdown hinders Fd/Fld reduction, leading to a bottleneck in MEP pathway flux. As such, our data are consistent with the conclusion that, in the absence of oxygen, Rnf uses the proton gradient to power the otherwise unfavorable transfer of electrons from NADH to Fd/Fld, which in turn drives essential processes such as isoprenoid biosynthesis (Fig. 4D).

Conversely, analysis of the F_1_F_O_ ATP synthase agrees with previous evidence suggesting the ATP synthase operates as a membrane potential generator (70–73). Partial knockdown of *atpC* resulted in lower membrane potential than an uninduced control (Fig. 4B). Furthermore, we reasoned that quantifying the ATP/ADP ratio for the *atpC* partial knockdown versus an uninduced control using metabolomics may illuminate the reaction direction. For example, if partial knockdown of *atp* genes results in an accumulation of ATP, this would suggest that, in the absence of knockdown, the F_1_F_O_ ATP synthase is functioning to hydrolyze ATP and pump ions outside the cytoplasm. Indeed, the ATP/ADP ratio increased relative to the non-targeting control for the two *atp* knockdowns with the clearest signal (Fig. 4E). This accumulation of ATP in the *atp* knockdown mutants indicates that during anaerobic growth in rich medium the F_1_F_O_ ATP synthase hydrolyzes ATP to generate PMF (Fig. 4F).

## Discussion

Alphaproteobacteria are fascinating models for fundamental biological processes as well as microbial powerhouses for industrial production of green energy. Despite this importance, core genes in Alphaproteobacteria are understudied due to the lack of both a simplified model genome and genetic tools capable of phenotyping all genes. This work advances our understanding of Alphaproteobacterial genes by establishing *Z. mobilis* as a streamlined, model microbe with a naturally reduced genome and employing a genome-scale, CRISPRi strategy for comprehensive phenotyping. We identified generally and conditionally essential genes that underpin the aerotolerant lifestyle of *Z. mobilis* and found broad conservation of these genes in other Alphaproteobacteria. Our analysis of Rnf and ATP synthase respiratory complexes highlights critical functions in maintaining/consuming PMF and points to a key role of Rnf in isoprenoid synthesis. Both our CRISPRi strategy and resulting insights into core gene functions are readily applicable to Alphaproteobacteria and beyond.

CRISPRi inducibility and titratability are invaluable for identifying conditional essentiality phenotypes. Because genes are instantly inactivated during Tn library construction, genes that are essential under the condition the library was constructed in are lost from the pool before downstream phenotyping experiments can begin. The only way to mitigate this issue would be to construct a new Tn library for every condition assayed—a considerable and impractical burden for investigators that are interested in probing many conditions to evaluate gene networks. CRISPRi inducibility mitigates this issue by separating library construction from fitness assays. As a result, we were able to define genes essential for aerobic growth despite constructing our CRISPRi library aerobically. An important example from our study is *recJ*, which has been defined as “essential” in previous Alphaproteobacterial Tn screens (63), but we show is conditionally essential during aerobic growth and dispensable anaerobically. Other genes that are canonically associated with oxygen stress (*e.g.*, *sod*) may also appear to be generally essential if Tn libraries are constructed aerobically. Titrating CRISPRi knockdowns allowed us to investigate the phenotypes of conditionally essential genes in the condition for which they are essential. This enabled us to define the mutator phenotype of *recJ* and PMF altering phenotypes of *rnf* and *atp* genes. Given the portability of CRISPRi (*e.g.*, Mobile-CRISPRi (74), we anticipate its broad utility in characterizing conditional essentiality.

We report that the broadly conserved respiratory complexes, Rnf and ATP synthase, are conditionally essential during *Z. mobilis* anaerobic growth and provide possible causes for their essentiality. Rnf (*<u>R</u>hodobacter* <u>n</u>itrogen fixation) complexes are widespread across prokaryotes and couple ion-motive force (Na^+^ or H^+^) with reversible Fd/Fld:NAD^+^ oxidoreductase activity (that is, oxidation of Fd_red_/Fld_red_ and reduction of NAD^+^, and vice versa) to accomplish diverse biological roles (67). In some anaerobes, Rnf is required for energy conservation: oxidation of Fd_red_/Fld_red_ is coupled with ion transport to establish ion-motive force necessary for ATP generation. Recently, the Rnf-ATP synthase supercomplex from *Thermatoga maritima* was purified and conclusively demonstrated to operate in this manner *in vitro* (75). In other organisms, ion transport occurs in the reverse direction, with ion-motive force powering the transfer of electrons from NADH to Fd/Fld. Canonically, Fd_red_/Fld_red_ then acts as a biologically powerful reductant and donates electrons to nitrogenase, a required step in N_2_ fixation (76, 77). *Z. mobilis* is known to fix N_2_ anaerobically in minimal medium lacking biologically available nitrogen (ammonium) (78). Under these conditions, *rnf* and other *nif* cluster genes associated with N_2_ fixation are upregulated accordingly (79), and their functions may be further regulated by protein phosphorylation status (80).

The present study extends the role of Rnf by demonstrating that Rnf is required for normal anaerobic growth and MEP pathway flux in *Z. mobilis*, even when ammonium is abundant. Here, using a multi-modal approach (genetic sensitivity to proton gradient disruption, ThT membrane potential assay, and metabolomics), we demonstrate that Rnf is a consumer of the proton gradient (Fig. 4), suggesting that it uses PMF to power reduction of Fd/Fld. Furthermore, we show that *rnfE* knockdown results in accumulation of MEP isoprenoid biosynthesis pathway intermediates (Fig. 4C). We suggest that MEP pathway flux is disrupted due to a deficiency in Fd_red_/Fld_red_, which donate electrons to the iron-sulfur cluster enzymes IspG and IspH in the MEP pathway (21, 81). Thus, this downstream disfunction of the essential MEP pathway may explain the essentiality of Rnf for anaerobic growth, in part or in full. Isoprenoids produced through the MEP pathway are biologically invaluable. In bacteria, isoprenoids act as electron carriers (quinones), pigments (carotenoids), membrane components (hopanoids), and signaling molecules (82). Given that isoprenoid biosynthesis is essential under all conditions for *Z. mobilis* (Table S9 and references) (46, 83, 84), the link between Rnf and MEP pathway flux during anaerobic growth suggests a separate mechanism may exist for generating Fd_red_/Fld_red_ to maintain flux during aerobic growth. Indeed, ferredoxin (*fdxA*; ZMO0220) and Fd/Fld reductase (*fpr*; ZMO1753) were both conditionally essential for aerobic growth in our screen. Given that *fpr* is also one of the most highly upregulated genes following oxygen exposure (21), we speculate that these genes may replenish Fd_red_/Fld_red_ aerobically, while Rnf performs this function anaerobically.

Ongoing engineering efforts aim to increase isoprenoid production by *Z. mobilis* for industrial use as therapeutics, food additives, fragrances, and sustainable biofuels (85). For example, Martien *et al.* observed that upregulation of *fpr*, *ispG*, and the iron-sulfur cluster assembly operon (*suf*) upon oxygen exposure coincided with improved flux through the MEP pathway, suggesting that manipulation of these genes could yield isoprenoid production strains with high MEP pathway flux (21). This proposal is supported by work in *E. coli*, where overexpression of Fpr, FldA, and MEP pathway enzymes increased flux through the MEP pathway (86). Furthermore, Khana *et al.* demonstrate that the iron-sulfur cluster enzymes IspG and IspH can modulate bottlenecks in the MEP pathway, and the authors point to electron-resupplying accessory proteins as possible engineering targets to further increase MEP pathway flux (81). The present work adds Rnf to this toolkit of potential engineering targets for enhancing isoprenoid production by *Z. mobilis*, especially for anaerobic industrial fermentations.

We also find that the F_1_F_O_ ATP synthase is conditionally essential for anaerobic growth. We further demonstrate that the F_1_F_O_ ATP synthase performs the essential function of hydrolyzing ATP to pump protons across the inner membrane and establish PMF. This role is consistent with previous conclusions surrounding *Z. mobilis* F_1_F_O_ ATP synthase function (70, 72, 73, 87). Our work both confirms this literature and additionally demonstrates that this function is essential for *Z. mobilis* anaerobic growth.

Our screen and CCCP sensitivity data also revealed an interesting contrast between the effects of *rnf* and *atp* knockdown on aerobic fitness. Specifically, while *rnf* knockdown appears to have no detectable impact on aerobic growth, knockdown of *atp* results in a mild fitness defect in the presence of oxygen (Fig. 4A, Table S9). This discrepancy could be explained by the possibly redundant function of *atp* with the *Z. mobilis* aerobic respiratory chain, which has an elusive function but may contribute to proton gradient generation and/or detoxification of oxygen (43, 87).

Given the clear connection between Rnf and F_1_F_O_ ATP synthase in *Thermatoga maritima* (75), it is tempting to speculate an equal-but-opposite model for *Z. mobilis* wherein the F_1_F_O_ ATP synthase establishes an ion gradient that is then directly used by Rnf. It is especially tempting given the long-anticipated discovery of a PMF-dissipating counterpart to F_1_F_O_ ATP synthase function (72, 87). Our data, however, point to additional biological complexity. If the main function of the F_1_F_O_ ATP synthase was to supply PMF for Rnf function, *atp* knockdown should yield a similar accumulation of MEP metabolites as occurs for *rnf* knockdown. However, we did not observe a consistent trend in MEP pathway metabolites for *atp* knockdown strains (Fig. S6). Multiple possible explanations exist. For example, disruption of the proton gradient by *atp* knockdown likely has widespread effects on cellular physiology (*e.g.,* in motility and transport) which may convolute biological outcomes, there may be additional ion translocators contributing to the proton gradient, and there may be other MEP pathway regulatory mechanisms at play (21, 81).

Herein, we establish *Z. mobilis* as a valuable genetic model for Alphaproteobacteria and deepen our understanding of how this naturally reduced genome bacterium adapts to the presence or absence of environmental oxygen. We anticipate that its use as a streamlined model with decreased genetic redundancy will simplify functional analysis of the basic biology of Alphaproteobacteria. We further expect that the novel biology gleaned from this study will aid in the development of prolific *Z. mobilis* strains for biofuel production.

## Materials and Methods

### Strains and growth conditions

Strains are listed in Table S1. Media recipes are in Table S2. *Escherichia coli* was grown in LB broth, Lennox (BD240230) at 37°C aerobically in a flask with shaking at 250 rpm, in a culture tube on a roller drum, or in a 96-well deep well plate with shaking at 900 rpm. *Zymomonas mobilis* was grown at 30°C aerobically or anaerobically (anaerobic chamber with 5% CO_2_, 5% H_2,_ balance N_2_) in RMG, ZRDM, or ZMM medium either statically in a culture tube or deep well plate, with stirring at 150 rpm, or in a Tecan Sunrise microplate reader statically with 60s shaking at high intensity prior to OD_600_ reads every 15 min. Media was solidified with 1.5-2% agar for growth on plates. Antibiotics were added when necessary: *E. coli* (100 µg/ml ampicillin (amp), 100 µg/ml carbenicillin (carb), 20 µg/ml chloramphenicol (cm) and *Z. mobilis* (100 µg/ml cm). Diaminopimelic acid (DAP) was added at 300 µM to support growth of *dap*-*E. coli* strains. Isopropyl β-D-1-thiogalactopyranoside (IPTG, 0.1-1 mM) was added where indicated. Strains were preserved in 15% glycerol at −80°C.

### General molecular biology techniques

Plasmids are listed in Table S3. Oligonucleotides are listed in Table S4. *pir*-dependent plasmids were propagated in *E. coli* strain BW25141 (sJMP146). Plasmids were purified using the GeneJet Plasmid Miniprep kit (Thermo K0503), the QIAprep Spin Miniprep Kit (Qiagen 27106), or the Purelink HiPure Plasmid Midiprep kit (Invitrogen K210005). Plasmids were digested with restriction enzymes from NEB and ligated using T4 DNA ligase (NEB M0202). DNA fragments were amplified using Q5 DNA polymerase (NEB 0491). PCR products were purified using the Monarch PCR & DNA Cleanup Kit (NEB T1030). Plasmids were transformed into electrocompetent *E. coli* cells using a BioRad Gene Pulser Xcell using the Ec1 program (0.1 cm cuvette, 1.80 kV, 1 pulse). Oligonucleotides were synthesized by Integrated DNA Technologies (Coralville, IA) or Agilent (Santa Clara, CA). Sequencing was performed by Functional Biosciences (Madison, WI) or the University of Wisconsin Biotechnology Center Next Generation Sequencing Core (Madison, WI).

### Alphaproteobacterial genome size analysis

Genome size analysis was performed using all complete National Center for Biotechnology Information (NCBI) genome entries for Alphaproteobacteria with a specific genus (*i.e.,* not “*Candidatus*,” “alpha,” or “uncultured”) as of July 7, 2021. The median number of genes (CDS) was calculated for each species and each genus.

### Comparative analysis of conserved and essential genes

OrthoFinder (88, 89) was used to identify orthologous groups among organisms. Annotated genomes of Alphaproteobacteria associated with published Tn-seq experiments (“Tractable Alphas”) obtained from the National Center for Biotechnology Information (NCBI) and corresponding published lists of essential genes for these organisms (35–40) were analyzed. The model organism *E. coli* served as a comparator (54, 55). See Table S5 for additional information on digital resources and links to custom scripts.

### *Z. mobilis* Mobile-CRISPRi individual gene and gene library construction

sgRNAs were designed to knock down all genes in *Z. mobilis* ZM4 using a custom Python script and GenBank CP023715.1 as detailed in reference (46). sgRNA-encoding sequences were cloned between the BsaI sites of Mobile-CRISPRi (MCi) plasmid pJMP2480. Methodology for cloning individual sgRNAs was described previously in detail (46, 47). Briefly, two 24-nucleotide (nt) oligonucleotides encoding an sgRNA were designed to overlap such that when annealed, their ends would be complementary to the BsaI-cut ends on the vector.

The pooled CRISPRi library was constructed by amplification of sgRNA-encoding spacer sequences (Table S6) from a custom pooled oligonucleotide library (SurePrint G7221A, Agilent) followed by ligation into the BsaI-digested MCi plasmid. Specifically, three pools of sgRNA-encoding inserts were generated by PCR amplification with primers oJMP197 and oJMP198 (Z1-genes), oJMP463 and oJMP464 (Z2-controls), and oJMP465 and oJMP466 (Z3-mismatches) from a 78-nt custom pooled oligonucleotide library with the following conditions per 300 µl reaction: 60 µl Q5 buffer, 9 µl GC enhancer, 6 µl 10mM each dNTPs, 15 µl each 10 µM forward and reverse primers, 6 µl 10 nM oligonucleotide library, 3 µl Q5 DNA polymerase, and 186 µl H_2_O with the following thermocycling parameters: 98°C, 30s; 15 cycles of: 98°C, 15s; 56°C, 15s; 72°C, 15s; 72°C, 10 min; 10°C, hold. Spin-purified PCR products were digested with BsaI-HF-v2 (R3733; NEB) and the size and integrity of full length and digested PCR products were confirmed on a 4% agarose e-gel (Thermo). The BsaI-digested PCR product (without further purification) was ligated into a BsaI-digested MCi plasmid as detailed in (47). The ligation was purified by spot dialysis on a nitrocellulose filter (Millipore VSWP02500) against 0.1 mM Tris, pH 8 buffer for 20 min prior to transformation by electroporation into *E. coli* strain BW25141 (sJMP146). Cells were plated at a density of ∼50,000 cells/plate on 150 mm LB-2% agar plates supplemented with carbenicillin. After incubation for 18 h at 37°C, colonies (∼1,300,000 (Z1-genes), 700,000 (Z2-controls), and 1,950,000 (Z3-mismatches) for ≥ 30 coverage/oligonucleotide) were scraped from the agar plates into LB, pooled, and the plasmid DNA was extracted from ∼1×10^11^ cells (10 ml at OD_600_ = 33 (Z1), 25 ml at OD_600_ = 11 (Z2), and 10 ml at OD_600_ = 36 (Z3) using a midiprep kit. This pooled Mobile-CRISPRi library was transformed by electroporation into *E. coli* mating strain sJMP3049 (20 ng plasmid DNA plus 90 µl electrocompetent cells, plated at a density of ∼30,000 cells/plate on 150 mm LB-2% agar plates supplemented with carbenicillin and DAP. After incubation for 18 h at 37°C, colonies (∼935,000 (Z1-genes), 124,000 (Z2-controls), and 247,000 (Z3-mismatches)) were scraped from the agar plates and pooled and resuspended in LB with DAP and 15% glycerol at OD_600_ (40 (Z1), 17 (Z2), and 34 (Z3)) and aliquots of the pooled CRISPRi libraries were stored as strains sJMP2618, 2619, and 2620 (Z1-genes, Z2-controls, and Z3-mismatches, respectively) at −80°C. Combined results from Z1-genes and Z2-controls are reported in this manuscript.

### Transfer of the Mobile-CRISPRi system to the *E. coli* or *Z. mobilis* chromosome

The MCi system was transferred to the Tn*7att* site on the chromosome of *Z. mobilis* by tri-parental conjugation of two donor strains—one with a mobilizable plasmid (pTn7C1) encoding Tn7 transposase and a second with a mobilizable plasmid containing a Tn7 transposon encoding the CRISPRi system—and the recipient strain *Z. mobilis* ZM4. All matings used the *E. coli* WM6026 donor strain, which is *pir^+^* to support pir-dependent plasmid replication, *dap^-^*, making it dependent on diaminopimelic acid (DAP) for growth, and encodes the RP4 transfer machinery required for conjugation. A detailed mating protocol for strains with individual sgRNAs was described previously (46, 47). Briefly, *E. coli* strains were grown ∼16 h and Z. *mobilis* strains were grown ∼24 h from single colonies. Cultures were spun at 4000 x *g* for 5 min and the cell pellets were washed twice with and equal volume of media (no antibiotic or DAP). For *E. coli* recipients, 100 µl of the washed donor and recipient strains were added to 700 µl LB, pelleted at ∼4000 x *g*, and the cells were placed on a nitrocellulose filter (Millipore HAWP02500) on an LB plate supplemented with DAP, and incubated at 37°C, ∼2 hr. For *Z. mobilis* recipients, 100 µl of the washed culture of donor strains and 500 µl of the recipient strain were added to 300 µl RMG, pelleted at ∼6000 x *g*, and the cells were placed on a nitrocellulose filter (Millipore HAWP02500) on an RMG plate supplemented with DAP, and incubated at 30°C, ∼24 h. Cells were removed from the filter by vortexing in 200 µl media, serially diluted, and grown with selection on LB-cm plates at 37°C (*E. coli*) or selection on RMG-cm plates at 30°C (*Z. mobilis*).

For pooled library construction, aliquots of the Tn*7* transposase donor strains (sJMP2618, 2619, and 2620) pooled library strains were thawed and diluted to OD_600_ = 10 in RMG. Overnight cultures of the *E. coli* Tn*7* transposase donor (sJMP2591) and *Z. mobilis* recipient strain (sJMP412) were spun down for 10 min at 4000 x *g* and the OD_600_ was normalized to 10. For each library, 2 ml of each strain was mixed and centrifuged for 10 min at 6000 x *g*. Pelleted cells were spotted on two RMG agar plates, and incubated for 39 h at 30°C prior to resuspension in RMG, serial dilution, and plating ∼40,000 CFU/150 mm RMG-cm plates solidified with 2% agar followed by incubation for 72 h at 30°C. Cells were scraped from plates and resuspended in RMG + 15% glycerol, the density was normalized to OD_600_ = 9 and aliquots were stored at - 80°C as strains sJMP2621, 2622, and 2623. Efficiency of trans-conjugation (colony forming units on RMG-cm vs. RMG) was ∼1 in 10^4^.

### Analysis of Mobile-CRISPRi reverse polarity in *Z. mobilis* or *E. coli* using a fluorescent reporter operon

An operon encoding mScarlet and sfGFP reporter genes was cloned into the Mobile-CRISPRi vector (pJMP2367) at the PmeI site. Nine sgRNAs per reporter gene were then designed and individually cloned into this plasmid vector, and the Mobile-CRISPRi constructs were transferred to *Z. mobilis* as described above.

For *Z. mobilis*, cultures from single colonies were grown in triplicate in 1 ml RMG-cm in a 96-well deep well plate covered with AeraSeal at 30°C without shaking for ∼24 h. *Z. mobilis* was subcultured 1:1000 inoculum:medium into two 96-well deep well plates containing either RMG or RMG + 1 mM IPTG and incubated as described above. The plates were centrifuged at 4000 x *g* for 10 min, the supernatant was removed, and the resulting cell pellets were resuspended in 1 ml PBS. Next, 200 µl of the cell suspension was transferred to a clear-bottom black microplate and measured in a Tecan Infinite microplate reader. Cells were shaken for 30s (linear amplitude: 2.5 mm) prior to measurement of cell density (OD_600_), GFP fluorescence (482/515 nm excitation/emission) and mScarlet fluorescence (560/605 nm). Experiments were repeated three times with 2-4 biological replicates each.

*E. coli* experiments were performed similarly, with the following modifications: LB was used in place of RMG, overnight growth was in 300 µl with shaking at 37°C for ∼18 h, *E. coli* was subcultured 1:10,000 inoculum:medium and grown 6-7 h, and *E. coli* cells were further diluted 1:2 in PBS (to OD_600_ ∼0.3-0.6) prior to measurement to avoid cell shadowing in dense culture.

OD-corrected fluorescence is reported relative to a no-sgRNA fluorescent control. One sgRNA (“S2,” which targets mScarlet) proved toxic to *Z. mobilis* and thus was excluded from analysis for this organism.

### Generation of predicted transcription units from RNA-seq data

We used previously published RNA-seq data (GSE139939) gathered from *Z. mobilis* ZM4 cultures grown to exponential and stationary phase in RMG under aerobic and anaerobic conditions to generate putative transcription unit (TU) assignments for *Z. mobilis* (11). Briefly, after adapter trimming with Cutadapt version 2.10 (90) and quality trimming with Trimmomatic version 0.39 (91), we aligned replicates from each condition and growth phase separately to the ZM4 reference chromosome (GenBank CP023715.1) and large plasmids (GenBanks CP023716.1, CP023717.1, CP023718.1, CP023719.1) using Rockhopper version 2.03 (92–94) to generate a set of predicted TUs for each condition and growth phase. Using custom Python scripts and Scipy version 1.5.2 (95), predicted transcript isoforms that covered the same genes for each growth phase were then parsed from Rockhopper output files and merged through single linkage hierarchical clustering using a pseudo-distance metric between two transcripts of “max size of compared transcripts in basepairs - number of basepairs that overlap”, and a cophenetic cutoff of 100 bp. For merged transcripts, the most extreme boundaries were used. TU assignments are included in Table S7.

### Library growth experiment

*Z. mobilis* CRISPRi libraries were incubated either in air or in an anaerobic chamber. The *Z. mobilis* CRISPRi libraries (sJMP2621, 2622, 2623) were revived by addition of 100 µl total frozen library stocks (OD_600_ = 9, 8:1:6 ratio Z1:Z2:Z3) into 100 ml RMG (starting OD_600_ = ∼0.01) in a 500 ml flask and incubated with stirring at 150 rpm at 30°C until OD_600_ = ∼0.2-0.3 (∼8 h) (initial timepoint = T*_i_*) in duplicate. These cultures were diluted back to OD_600_ = 0.01 in RMG + 1 mM IPTG in duplicate and incubated until OD_600_ = ∼0.2-0.3 (∼8 h) (final timepoint = T*_f_*), at which point they were again diluted back to OD_600_ = 0.01 in RMG + 1 mM IPTG in duplicate and incubated until OD_600_ = ∼0.2-0.3 (∼8 h). Cells were pelleted at 6000 x *g* from 30 ml (T*_i_*) or 10 ml (T*_f_*), washed with 1X PBS and stored at −20°C for DNA extraction. Aerobic and anaerobic experiments were each done twice on separate days.

### Sequencing library samples

DNA was extracted from cell pellets with the DNeasy gDNA extraction kit (Qiagen) according to the manufacturer’s protocol, resuspending in a final volume of 100 µl with an average yield of ∼50 ng/µl. The sgRNA-encoding region was amplified using Q5 DNA polymerase (NEB) in a 100 µl reaction with 2 µl gDNA (∼100 ng) and primers oJMP697 and oJMP698 (nested primers with partial adapters for index PCR with Illumina TruSeq adapter) according to the manufacturer’s protocol using a BioRad C1000 thermal cycler with the following program: 98°C, 30s then 16 cycles of: 98°C, 15s; 65°C, 15s; 72°C, 15s. PCR products were spin purified and eluted in a final volume of 20 µl for a final concentration of ∼20 ng/µl). Samples were sequenced by the UW-Madison Biotech Center Next Generation Sequencing Core facility. Briefly, PCR products were amplified with nested primers containing i5 and i7 indexes and Illumina TruSeq adapters followed by bead cleanup, quantification, pooling and running on a NovaSeq 6000 (150 bp paired end reads).

### Counting sgRNA sequences

sgRNA-encoding spacer sequences were counted using the seal.sh script from the BBTools package (Release: March 28, 2018). Briefly, paired FASTQ files from amplicon sequencing were aligned in parallel to a reference file that contained the spacer sequences cloned into the library. Alignment was performed using *k*-mers of 20 nucleotide length—equal to the length of the spacer sequence. For more information on digital resources and links to custom scripts, see Table S5.

### Comparisons between conditions

Log_2_ fold change (LFC) and confidence intervals were computed using edgeR. Briefly, trended dispersion of sgRNA-encoding spacers was estimated and imputed into a quasi-likelihood negative binomial log-linear model. Changes in abundance and the corresponding false discovery rates (FDRs) were computed for each spacer in each condition. Finally, gene-level fitness scores were obtained by calculating the median LFC of the spacers targeting each gene; gene-level significance was calculated by computing the Stouffer’s *p*-value (poolr R package) using the FDRs of the spacers targeting each gene. For more information on digital resources and links to custom scripts, see Table S5.

### Classifying gene essentiality and fitness defects

Genes with significance ≤ 0.05 and median LFC ≤ −3 under aerobic and anaerobic conditions were considered generally essential. Genes with significance ≤ 0.05 and median LFC ≤ −3 under only one condition were classified as conditionally essential. Genes with significance ≤ 0.05 and median LFC ≤ −1 in one or both conditions were classified as having a fitness defect in the respective condition(s).

### Analysis of individual CRISPRi knockdown strains for essentiality using spot dilution assays

Phenotypes of gene knockdowns identified in the pooled screen were assayed by constructing individual *Z. mobilis* strains with sgRNAs targeting 31 putative generally essential genes, 20 putative aerobic essential or aerobic-specific fitness defect genes, 24 putative anaerobic essential or anaerobic-specific fitness defect genes, and 6 non-targeting control sgRNA strains. Individual colonies from each strain were picked in duplicate (2 biological replicates) into 1 ml RMG-cm into 96-well deep well plates, covered with AeraSeal, and incubated at 30°C for 24 h to saturation without shaking. Cultures were serially (1:10) diluted in RMG in a sterile V-bottom 96-well microplate and spotted onto RMG or RMG + 1 mM IPTG agar plates (OmniTray, Thermo 242811, 33 ml 2% agar medium) using a 96-well manual pinning tool (V&P Scientific; VP407A pin tool, VP381N microplate aligner, VP380 OmniTray agar plate aligner) in triplicate (3 technical replicates) and incubated either aerobically or anaerobically at 30°C for 72 h. Growth was measured using a fitness score (number of serial dilutions resulting in a visible spot) and colony size score (small, large, or control-like). Growth was scored by 3 individuals; the median fitness score and the most prevalent colony size score are reported.

### Analysis of *recJ* CRISPRi knockdown strains mutation rate in the presence of rifampicin

Acquisition of rifampicin resistance by a *Z. mobilis recJ* knockdown strain (sJMP10367) compared to a non-targeting control strain (sJMP10366) and parent strain (sJMP10365) was assayed as follows: ∼2.4 x 10^6^ cells resuspended off plates into RMG (200 µl of a 10^-5^ dilution of cells normalized to OD_600_ = 9) were plated in triplicate on RMG + 20 µM IPTG and incubated at 30°C for 90 h to induce partial knockdown. ∼2.6 x 10^8^ cells from each replicate resuspended off plates into RMG (200 µl of cells normalized to OD_600_ = 12) were plated in quadruplicate on RMG or RMG + 4 µM rifampicin (rif) and incubated aerobically, and on RMG or RMG + 8 µM rif and incubated anaerobically, at 30°C in the dark for 96 h. Colony forming units (CFU) were compared for growth +/− rif.

### Fitness measurement of *Z. mobilis atp* and *rnf* partial knockdowns with proton motive force disruption by CCCP

Fitness challenge experiments were performed in clear 96-well microplates inoculated 1:50 (inoculum:medium) in duplicate from a saturated 5-ml overnight culture grown in RMG under either aerobic or anaerobic conditions. For anaerobic experiments, the 96-well microplate was placed in the anaerobic chamber overnight prior to inoculation. *atp* knockdown (sJMP6104, sJMP6105, sJMP6109), *rnf* knockdown (sJMP6103), and non-targeting CRISPRi (sJMP6101) strains were grown in RMG with sublethal concentrations of inducer (50 μM IPTG), carbonyl cyanide 3-chlorophenylhydrazone (CCCP) (8 μg/ml, from stock dissolved in DMSO), and/or DMSO (0.5%) as a control. Growth data were collected in a Tecan Sunrise plate reader for ∼48 h. Experiments were repeated three times with two biological replicates each.

Actual relative fitness was measured by calculating the empirical area under the curve (auc_e) for each growth curve using the Growthcurver R package (version 0.3.1) relative to the DMSO-only control for each strain. Following the multiplicative theory for synergy (49), predicted relative fitness for a combination of two conditions was calculated by multiplying the actual relative fitness values obtained from each condition individually (*i.e.,* induction and CCCP). Actual and predicted fitness values are reported relative to those of the non-targeting control strain.

### Detection of membrane potential using Thioflavin T fluorescence

Overnight cultures of *rnfE* knockdown (MT271), *atpC* knockdown (MT272) and a non-targeting CRISPRi strain (MT270) were inoculated from single colonies in ZRDM + 70 μg/ml cm in duplicate and grown at 30°C in an anaerobic chamber. Cultures were diluted in fresh ZRDM-cm + 10 μM Thioflavin T (ThT) to OD_600_ = 0.04. 50 μM IPTG inducer was added as indicated. 200 μl from each biological replicate was transferred to a 96-well microplate (flat, black, clear-bottom, Greiner) in triplicate. Cells were grown at 30°C with shaking in a BioTek Synergy HTX Multimode Reader in an anaerobic chamber. OD_600_ and ThT fluorescence (optical filters 460/40 nm and 528/20 nm for excitation and emission, respectively) were measured every 15 minutes for 30 h. Average ThT fluorescence and OD_600_ values were corrected by subtracting background values from abiotic controls along the time course.

### *Z. mobilis* growth curves in RMG or ZRDM

Aerobic cultures grown overnight in 5 ml RMG were washed twice in ZMM, then resuspended in ZMM to original volume. The washed cultures were diluted 1:50 (2 µl into 98 µl) into growth medium (RMG or ZRDM) in a 96-well microplate and grown ∼24 hours in a Tecan Sunrise microplate reader in the anaerobic chamber. Growth curves represent 4-6 biological replicates across 2-3 experiments.

### Liquid chromatography-mass spectrometry metabolomics for *atp* and *rnf* knockdown strains

*Z. mobilis* was grown at 30°C in an anaerobic chamber. *Z. mobilis* cultures of *atp* knockdowns (sJMP6104, sJMP6105, sJMP6109), *rnfE* knockdown (sJMP6103) and non-targeting CRISPRi (sJMP6101) strains were grown overnight, stationary, in 10 ml RMG in test tubes from single colonies, diluted ∼1:1000 into 60 ml RMG in 125-ml flasks and grown with stirring (120 rpm) until OD_600_ ∼0.7-0.9 to use as inoculum for experimental flasks. Experimental flasks were inoculated to OD_600_ = 0.05 in 100 ml ZRDM + 50 µM IPTG in 250-ml flasks with stirring. From this flask, 30 ml was transferred into each of three separate 125-ml flasks with stirring. When these cultures reached OD = 0.5, 10 ml from each experimental flask was harvested by vacuum filtration onto a 0.45 µm nylon membrane filter (47 mm diameter). Cells were resuspended off the filter into 1.5 ml ice cold solvent (40:40:20 acetonitrile:methanol:water), and the solvent was stored in a microcentrifuge tube at −80°C until ready for same-day processing.

As an internal standard, wild-type *Z. mobilis* ZM4 was grown in ZMM + labeled glucose (Table S6). Wild-type *Z. mobilis* was prepared as described above, with the following modifications: overnight RMG cultures were diluted into 10 ml ZMM + labeled glucose, a single experimental flask containing 30 ml ZMM + labeled glucose was inoculated, and 10 ml was collected in duplicate from the experimental flask.

Collected samples were spun in a microcentrifuge at max speed for 10 min at 4°C. Each sample’s supernatant was mixed 1:1 with supernatant from the labeled glucose internal standard and dried under a nitrogen gas stream.

Metabolomics LC-MS analyses were conducted utilizing a Vanquish ultra-high-performance liquid chromatography (UHPLC) system (Thermo Scientific), coupled with a Q Exactive™ hybrid quadrupole-Orbitrap™ mass spectrometer (Thermo Scientific) using electrospray ionization in negative-ion mode. The chromatography was performed at 25°C with a reverse-phase C18 column of 2.1 × 100 mm with 1.7 μm particle size (Water™ from Acquity UHPLC BEH). Solvent A (97:3 H_2_O:methanol with 10 mM tributylamine adjusted to pH 8.2 using 10 mM acetic acid) and Solvent B (100% methanol) were used in a gradient manner: 0-2.5 min with 5% B, 2.5-17 min with a linear gradient from 5% B to 95% B, 17-19.5 min with 95% B, 19.5-20 min with a linear gradient from 95% B to 5% B, and 20-25 min with 5% B. The flow rate was constant at 0.2 ml/min. For targeted metabolomics, the eluent was injected into the MS for analysis until 18 min, after which the flow was redirected to waste. The MS parameters included full MS-SIM (single ion monitoring) scanning between 70 and 1,000 m/z and 160 and 815 m/z for the targeted metabolomics and MEP metabolite-specific methods, respectively. The automatic control gain (ACG) target was 1e6, with a maximum injection time (IT) of 40 ms and a resolution of 70,000 full width at half maximum (FWHM).

## Acknowledgements

This work was supported by the U.S. Department of Energy, Office of Science, Office of Biological and Environmental Research, Great Lakes Bioenergy Research Center under Award Number DE-SC0018409. A.L.E. was supported by the Biotechnology Training Program (NIH 5T32GM135066) and a GRFP from the NSF. R.D.W. was supported by the Predoctoral Training Program in Genetics (NIH 5T32GM007133). We thank members of the Great Lakes Bioenergy Research Center, especially the Peters, Kiley, Pfleger, and Sato labs for helpful feedback. We thank Piyush Lal and Tricia Kiley for strains and plasmids as well as Lisa Liu, Trey Sato, and the University of Wisconsin Biotechnology Center for technical assistance. PH-PLCD1_mScarlet_IRES_SYFP2_PH_N1 was a gift from Dorus Gadella (Addgene plasmid # 85069 (96).

---

**Figure S1.**
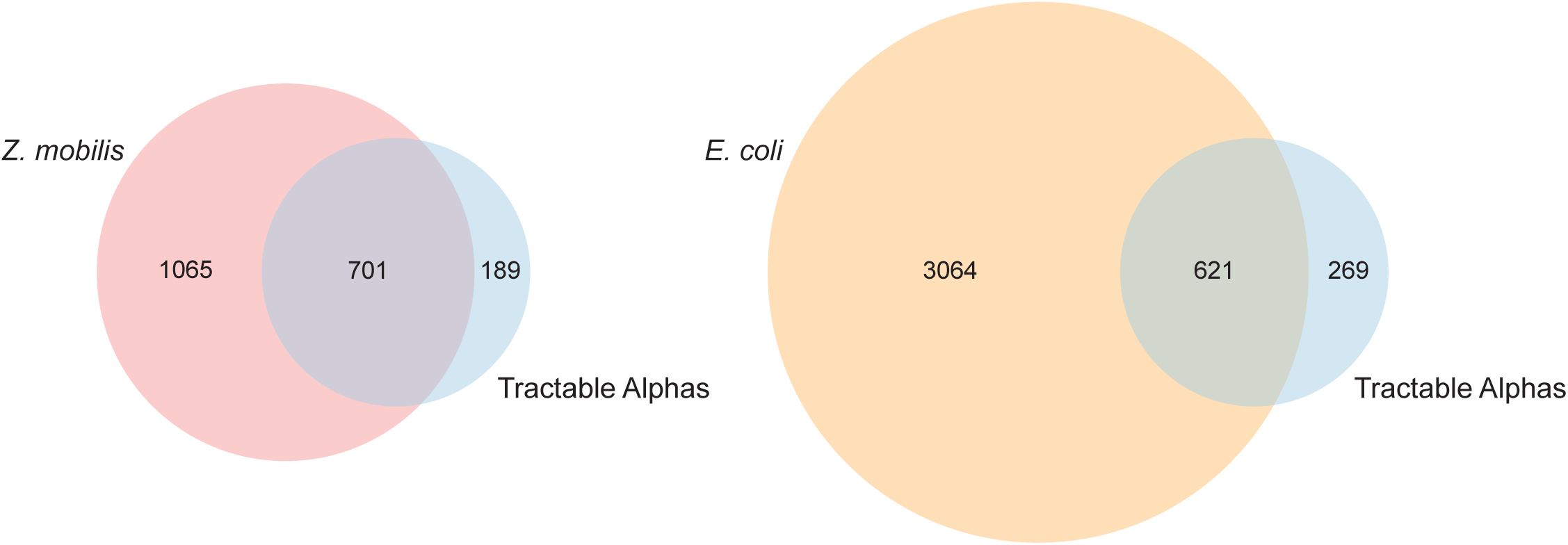
Gene homology of *Z. mobilis* or *E. coli* with other Alphaproteobacteria. Blue overlapping circles represent protein-coding orthologs shared across all seven Tractable Alphas (*C. crescentus, B. subvibrioides, A. tumefaciens, B. diazoefficiens, R. palustris, R. sphaeroides, S. wittichii*). Red and orange overlapping circles represent all protein-coding orthologs in *Z. mobilis* and *E. coli*, respectively.

**Figure S2.**
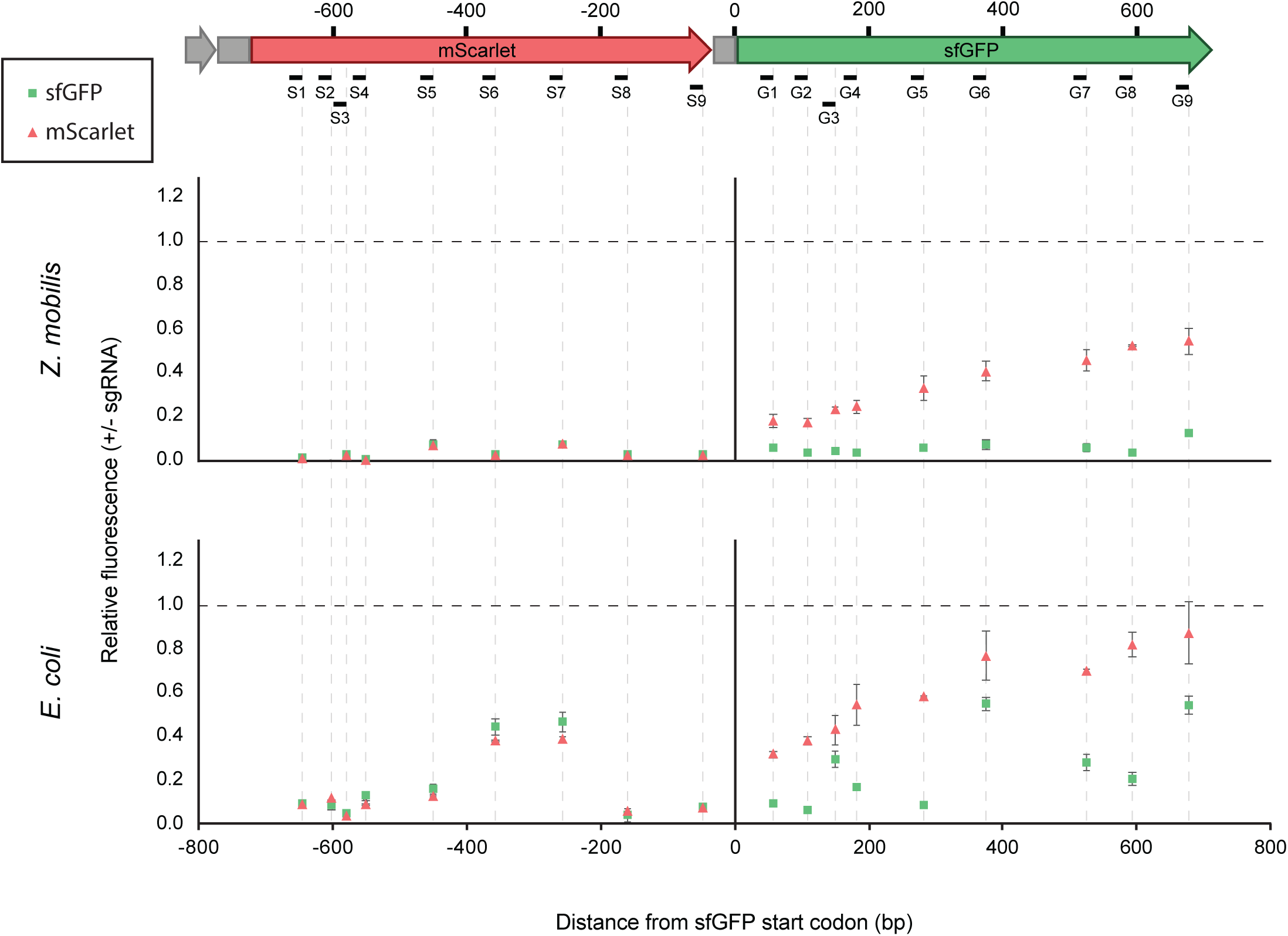
Reverse polarity in *Z. mobilis* CRISPRi. (A) Fluorescent reporter operon structure. sgRNAs targeting *mScarlet* (S1-9) and *sfGFP* (G1-9) are noted. (B-C) Fluorescence of mScarlet (pink triangle) and sfGFP (green square) for corresponding knockdown mutants in (B) *Z. mobilis* and (C) *E. coli*. Fluorescence is reported relative to a non-targeting sgRNA control (horizontal dashed line). Data represent three experiments with 2-4 biological replicates each.

**Figure S3.**
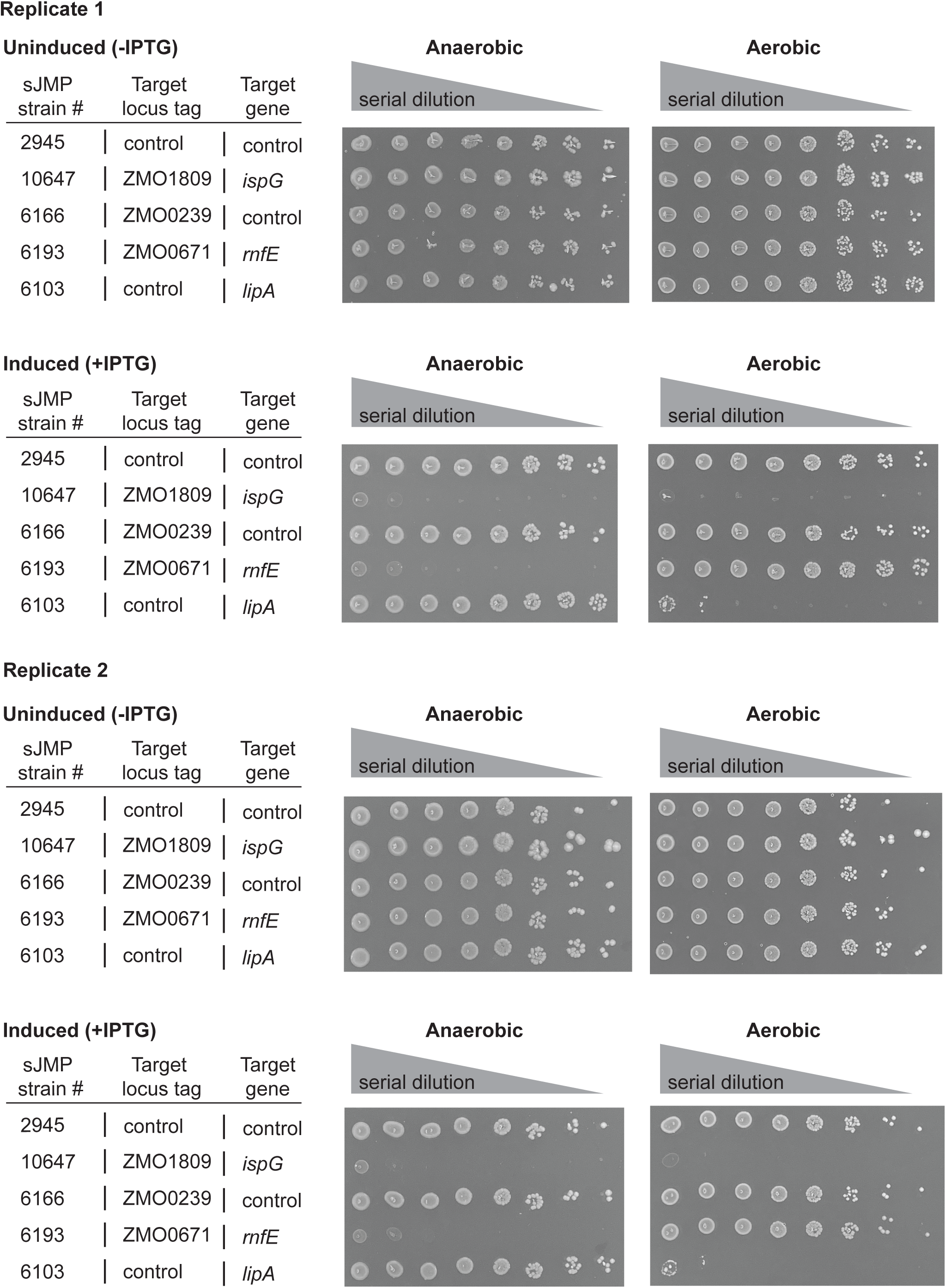
Example verification of *Z. mobilis* CRISPRi library screen phenotypes for generally essential (*ispG*), anaerobic essential (*rnfE*), and aerobic essential (*lipA*) genes using spot plates (see Methods).

**Figure S4.**
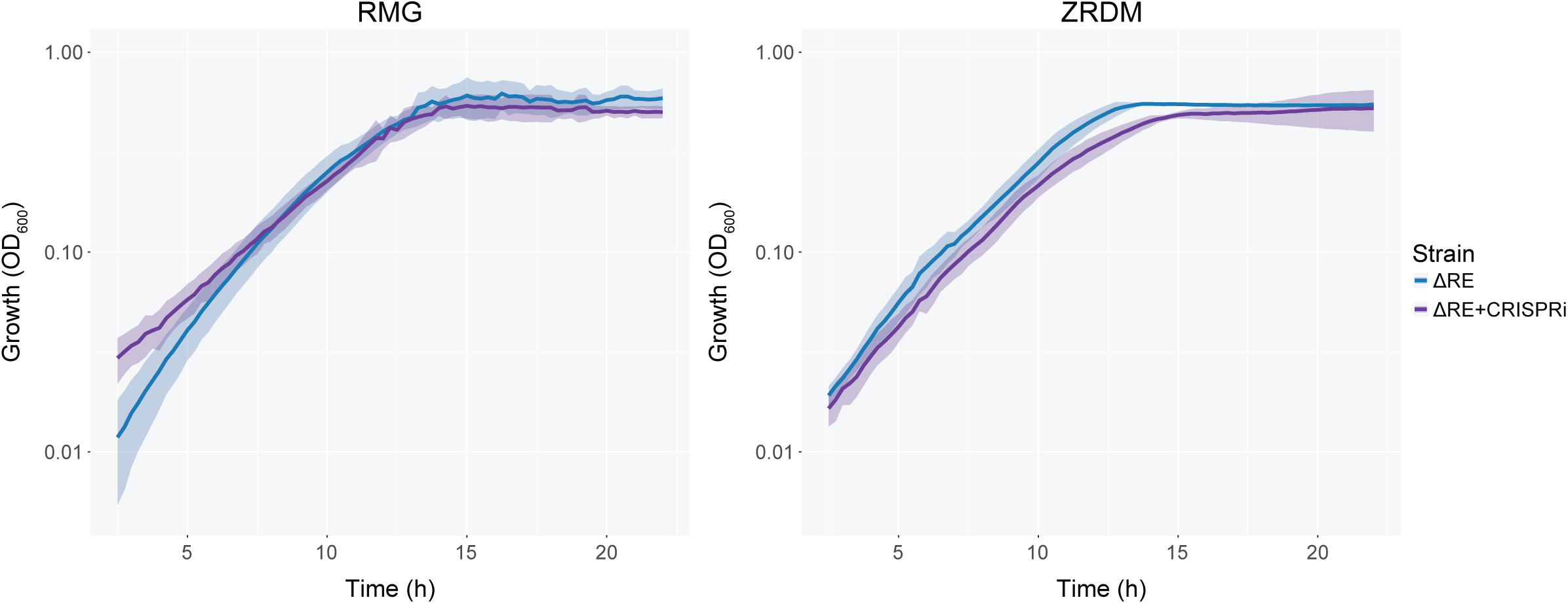
Growth of *Z. mobilis* strains in Rich Medium with Glucose (RMG) or *Zymomonas* Rich Defined Medium (ZRDM) without induction. ΔRE, restriction-deficient parent strain (sJMP412); ΔRE+CRISPRi, ΔRE strain with non-targeting Mobile-CRISPRi (sJMP2554). Shaded ribbons represent one standard deviation from the mean. Data represent 4-6 replicates across 2-3 experiments.

**Figure S5.**
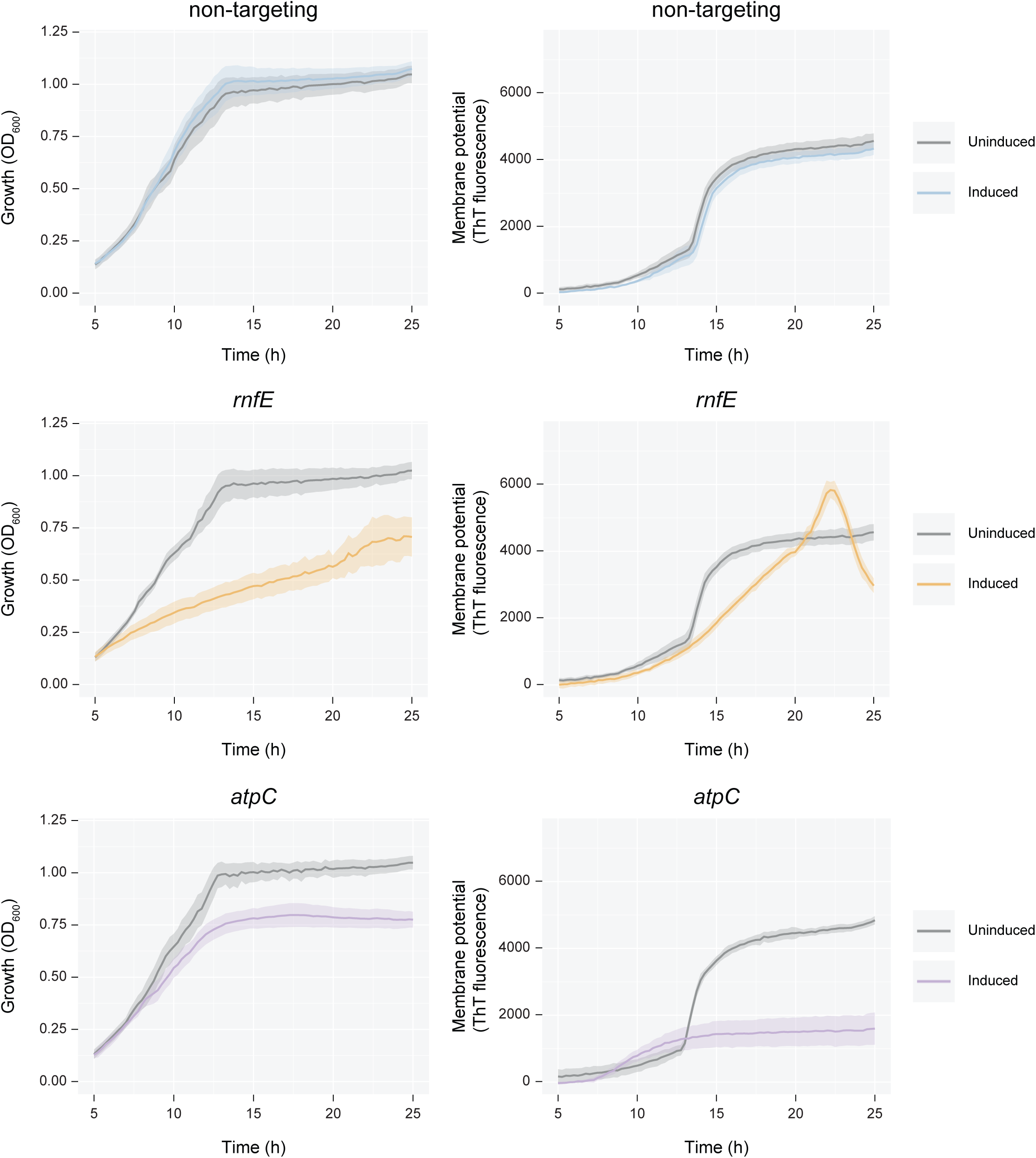
Growth (left column) and membrane potential (right column) for strains corresponding to Figure 4B. Shaded ribbons represent one standard deviation from the mean. Data represent six replicates.

**Figure S6.**
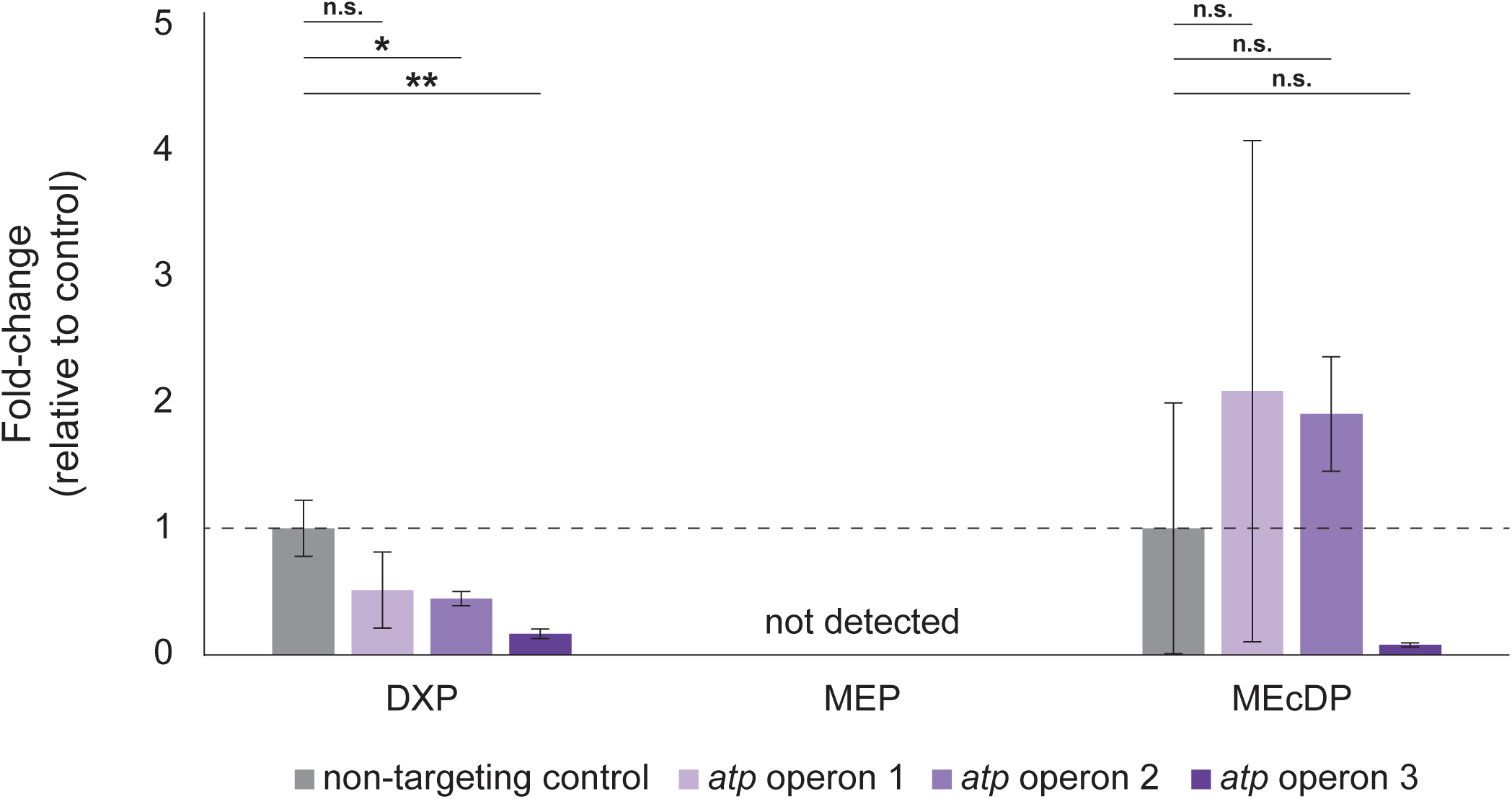
Mass spectroscopy metabolomics measurement of MEP pathway intermediates for (leftmost gray bars and dashed line) non-targeting CRISPRi control and (righthand purple bars) *atp* partial knockdown. n.s., not significant; *, *p* < 0.05; **, *p* < 0.01 by two-tailed Student’s T-test. *atp* operon 1 knockdown targets *atpA*; *atp* operon 2 knockdown targets *atpC*; *atp* operon 3 knockdown targets *atpF*. DXP, 1-deoxy-D-xylulose 5-phosphate; MEP, 2-C-methyl-D-erythritol 4-phosphate; MEcDP, 2-C-methyl-D-erythritol-2,4-cyclodiphosphate. Data represent three replicates.

## References

1. LeBlanc N, Charles TC. 2022. Bacterial genome reductions: Tools, applications, and challenges. Front Genome Ed 4:957289.

2. Choe D, Cho S, Kim SC, Cho B-K. 2016. Minimal genome: Worthwhile or worthless efforts toward being smaller? Biotechnol J 11:199–211.

3. Bu Q-T, Yu P, Wang J, Li Z-Y, Chen X-A, Mao X-M, Li Y-Q. 2019. Rational construction of genome-reduced and high-efficient industrial Streptomyces chassis based on multiple comparative genomic approaches. Microb Cell Fact 18:16.

4. Csörgo B, Fehér T, Tímár E, Blattner FR, Pósfai G. 2012. Low-mutation-rate, reduced-genome Escherichia coli: an improved host for faithful maintenance of engineered genetic constructs. Microb Cell Fact 11:11.

5. Qiao W, Liu F, Wan X, Qiao Y, Li R, Wu Z, Saris PEJ, Xu H, Qiao M. 2021. Genomic Features and Construction of Streamlined Genome Chassis of Nisin Z Producer Lactococcus lactis N8. Microorganisms 10:47.

6. Pósfai G, Plunkett G, Fehér T, Frisch D, Keil GM, Umenhoffer K, Kolisnychenko V, Stahl B, Sharma SS, de Arruda M, Burland V, Harcum SW, Blattner FR. 2006. Emergent properties of reduced-genome Escherichia coli. Science 312:1044–1046.

7. Liu Y, Feng J, Pan H, Zhang X, Zhang Y. 2022. Genetically engineered bacterium: Principles, practices, and prospects. Front Microbiol 13:997587.

8. Hashimoto M, Ichimura T, Mizoguchi H, Tanaka K, Fujimitsu K, Keyamura K, Ote T, Yamakawa T, Yamazaki Y, Mori H, Katayama T, Kato J. 2005. Cell size and nucleoid organization of engineered Escherichia coli cells with a reduced genome. Mol Microbiol 55:137–149.

9. Kurokawa M, Ying B-W. 2019. Experimental Challenges for Reduced Genomes: The Cell Model Escherichia coli. Microorganisms 8:3.

10. Iwadate Y, Honda H, Sato H, Hashimoto M, Kato J. 2011. Oxidative stress sensitivity of engineered Escherichia coli cells with a reduced genome. FEMS Microbiol Lett 322:25–33.

11. Vera JM, Ghosh IN, Zhang Y, Hebert AS, Coon JJ, Landick R. 2020. Genome-Scale Transcription-Translation Mapping Reveals Features of Zymomonas mobilis Transcription Units and Promoters. mSystems 5.

12. Yang S, Vera JM, Grass J, Savvakis G, Moskvin OV, Yang Y, McIlwain SJ, Lyu Y, Zinonos I, Hebert AS, Coon JJ, Bates DM, Sato TK, Brown SD, Himmel ME, Zhang M, Landick R, Pappas KM, Zhang Y. 2018. Complete genome sequence and the expression pattern of plasmids of the model ethanologen Zymomonas mobilis ZM4 and its xylose-utilizing derivatives 8b and 2032. Biotechnology for Biofuels 11:125.

13. He MX, Wu B, Qin H, Ruan ZY, Tan FR, Wang JL, Shui ZX, Dai LC, Zhu QL, Pan K, Tang XY, Wang WG, Hu QC. 2014. Zymomonas mobilis: a novel platform for future biorefineries. Biotechnol Biofuels 7:101.

14. Qiu M, Shen W, Yan X, He Q, Cai D, Chen S, Wei H, Knoshaug EP, Zhang M, Himmel ME, Yang S. 2020. Metabolic engineering of Zymomonas mobilis for anaerobic isobutanol production. Biotechnology for Biofuels 13:15.

15. Ong WK, Courtney DK, Pan S, Andrade RB, Kiley PJ, Pfleger BF, Reed JL. 2020. Model-driven analysis of mutant fitness experiments improves genome-scale metabolic models of Zymomonas mobilis ZM4. PLOS Computational Biology 16:e1008137.

16. Felczak MM, Bowers RM, Woyke T, TerAvest MA. 2021. Zymomonas diversity and potential for biofuel production. Biotechnol Biofuels 14:112.

17. Glass JI, Assad-Garcia N, Alperovich N, Yooseph S, Lewis MR, Maruf M, Hutchison CA, Smith HO, Venter JC. 2006. Essential genes of a minimal bacterium. Proc Natl Acad Sci U S A 103:425–430.

18. Rogers PL, Jeon YJ, Lee KJ, Lawford HG. 2007. Zymomonas mobilis for fuel ethanol and higher value products. Adv Biochem Eng Biotechnol 108:263–288.

19. Franden MA, Pilath HM, Mohagheghi A, Pienkos PT, Zhang M. 2013. Inhibition of growth of Zymomonas mobilis by model compounds found in lignocellulosic hydrolysates. Biotechnol Biofuels 6:99.

20. Hu M, Chen X, Huang J, Du J, Li M, Yang S. 2021. Revitalizing the ethanologenic bacterium Zymomonas mobilis for sugar reduction in high-sugar-content fruits and commercial products. Bioresour Bioprocess 8:119.

21. Martien JI, Hebert AS, Stevenson DM, Regner MR, Khana DB, Coon JJ, Amador-Noguez D. Systems-Level Analysis of Oxygen Exposure in Zymomonas mobilis: Implications for Isoprenoid Production. mSystems 4:e00284–18.

22. Goodman AL, Wu M, Gordon JI. 2011. Identifying microbial fitness determinants by insertion sequencing using genome-wide transposon mutant libraries. Nat Protoc 6:1969–1980.

23. van Opijnen T, Bodi KL, Camilli A. 2009. Tn-seq: high-throughput parallel sequencing for fitness and genetic interaction studies in microorganisms. Nat Methods 6:767–772.

24. Skerker JM, Leon D, Price MN, Mar JS, Tarjan DR, Wetmore KM, Deutschbauer AM, Baumohl JK, Bauer S, Ibáñez AB, Mitchell VD, Wu CH, Hu P, Hazen T, Arkin AP. 2013. Dissecting a complex chemical stress: chemogenomic profiling of plant hydrolysates. Molecular Systems Biology 9:674.

25. Brenac L, Baidoo EEK, Keasling JD, Budin I. 2019. Distinct functional roles for hopanoid composition in the chemical tolerance of Zymomonas mobilis. Molecular Microbiology 112:1564–1575.

26. Fuchino K, Wasser D, Soppa J. 2021. Genome Copy Number Quantification Revealed That the Ethanologenic Alpha-Proteobacterium Zymomonas mobilis Is Polyploid. Front Microbiol 12:705895.

27. Fuchino K, Chan H, Hwang LC, Bruheim P. 2021. The Ethanologenic Bacterium Zymomonas mobilis Divides Asymmetrically and Exhibits Heterogeneity in DNA Content. Appl Environ Microbiol 87:e02441–20.

28. Banta AB, Enright AL, Siletti C, Peters JM. 2020. A High-efficacy CRISPRi System for Gene Function Discovery in Zymomonas mobilis. bioRxiv 2020.07.06.190827.

29. Qi LS, Larson MH, Gilbert LA, Doudna JA, Weissman JS, Arkin AP, Lim WA. 2013. Repurposing CRISPR as an RNA-guided platform for sequence-specific control of gene expression. Cell 152:1173–1183.

30. Hawkins JS, Silvis MR, Koo B-M, Peters JM, Osadnik H, Jost M, Hearne CC, Weissman JS, Todor H, Gross CA. 2020. Mismatch-CRISPRi Reveals the Co-varying Expression-Fitness Relationships of Essential Genes in Escherichia coli and Bacillus subtilis. Cell Syst https://doi.org/10.1016/j.cels.2020.09.009.

31. Lee JS, Jin SJ, Kang HS. 2001. Molecular organization of the ribosomal RNA transcription unit and the phylogenetic study of Zymomonas mobilis ZM4. Mol Cells 11:68–74.

32. Seo J-S, Chong H, Park HS, Yoon K-O, Jung C, Kim JJ, Hong JH, Kim H, Kim J-H, Kil J-I, Park CJ, Oh H-M, Lee J-S, Jin S-J, Um H-W, Lee H-J, Oh S-J, Kim JY, Kang HL, Lee SY, Lee KJ, Kang HS. 2005. The genome sequence of the ethanologenic bacterium Zymomonas mobilis ZM4. Nat Biotechnol 23:63–68.

33. Serbus LR, Casper-Lindley C, Landmann F, Sullivan W. 2008. The genetics and cell biology of Wolbachia-host interactions. Annu Rev Genet 42:683–707.

34. Helminiak L, Mishra S, Kim HK. 2022. Pathogenicity and virulence of Rickettsia. Virulence 13:1752–1771.

35. Burger BT, Imam S, Scarborough MJ, Noguera DR, Donohue TJ. 2017. Combining Genome-Scale Experimental and Computational Methods To Identify Essential Genes in Rhodobacter sphaeroides. mSystems 2:e00015–17.

36. Pechter KB, Gallagher L, Pyles H, Manoil CS, Harwood CS. 2015. Essential Genome of the Metabolically Versatile Alphaproteobacterium Rhodopseudomonas palustris. J Bacteriol 198:867–876.

37. Curtis PD, Brun YV. 2014. Identification of essential alphaproteobacterial genes reveals operational variability in conserved developmental and cell cycle systems. Mol Microbiol 93:713–735.

38. Christen B, Abeliuk E, Collier JM, Kalogeraki VS, Passarelli B, Coller JA, Fero MJ, McAdams HH, Shapiro L. 2011. The essential genome of a bacterium. Mol Syst Biol 7:528.

39. Roggo C, Coronado E, Moreno-Forero SK, Harshman K, Weber J, van der Meer JR. 2013. Genome-wide transposon insertion scanning of environmental survival functions in the polycyclic aromatic hydrocarbon degrading bacterium Sphingomonas wittichii RW1. Environ Microbiol 15:2681–2695.

40. Baraquet C, Dai W, Mendiola J, Pechter K, Harwood CS. 2021. Transposon sequencing analysis of Bradyrhizobium diazoefficiens 110spc4. Sci Rep 11:13211.

41. Sootsuwan K, Lertwattanasakul N, Thanonkeo P, Matsushita K, Yamada M. 2008. Analysis of the respiratory chain in Ethanologenic Zymomonas mobilis with a cyanide-resistant bd-type ubiquinol oxidase as the only terminal oxidase and its possible physiological roles. J Mol Microbiol Biotechnol 14:163–175.

42. Balodite E, Strazdina I, Galinina N, McLean S, Rutkis R, Poole RK, Kalnenieks U. 2014. Structure of the Zymomonas mobilis respiratory chain: oxygen affinity of electron transport and the role of cytochrome c peroxidase. Microbiology (Reading) 160:2045–2052.

43. Felczak MM, TerAvest MA. 2023. Respiration is essential for aerobic growth of Zymomonas mobilis ZM4. bioRxiv https://doi.org/10.1101/2023.03.02.530925.

44. Jacobson TB, Adamczyk PA, Stevenson DM, Regner M, Ralph J, Reed JL, Amador-Noguez D. 2019. 2H and 13C metabolic flux analysis elucidates in vivo thermodynamics of the ED pathway in Zymomonas mobilis. Metabolic Engineering 54:301–316.

45. Lee KY, Park JM, Kim TY, Yun H, Lee SY. 2010. The genome-scale metabolic network analysis of Zymomonas mobilis ZM4 explains physiological features and suggests ethanol and succinic acid production strategies. Microbial Cell Factories 9:94.

46. Banta AB, Enright AL, Siletti C, Peters JM. 2020. A High-efficacy CRISPRi System for Gene Function Discovery in Zymomonas mobilis. Appl Environ Microbiol https://doi.org/10.1128/AEM.01621-20.

47. Banta AB, Ward RD, Tran JS, Bacon EE, Peters JM. 2020. Programmable Gene Knockdown in Diverse Bacteria Using Mobile-CRISPRi. Curr Protoc Microbiol 59:e130.

48. Cui L, Vigouroux A, Rousset F, Varet H, Khanna V, Bikard D. 2018. A CRISPRi screen in E. coli reveals sequence-specific toxicity of dCas9. Nat Commun 9:1912.

49. Peters JM, Colavin A, Shi H, Czarny TL, Larson MH, Wong S, Hawkins JS, Lu CHS, Koo B- M, Marta E, Shiver AL, Whitehead EH, Weissman JS, Brown ED, Qi LS, Huang KC, Gross CA. 2016. A Comprehensive, CRISPR-based Functional Analysis of Essential Genes in Bacteria. Cell 165:1493–1506.

50. Peters JM, Silvis MR, Zhao D, Hawkins JS, Gross CA, Qi LS. 2015. Bacterial CRISPR: accomplishments and prospects. Curr Opin Microbiol 27:121–126.

51. Zhang R, Xu W, Shao S, Wang Q. 2021. Gene Silencing Through CRISPR Interference in Bacteria: Current Advances and Future Prospects. Front Microbiol 12:635227.

52. Rock JM, Hopkins FF, Chavez A, Diallo M, Chase MR, Gerrick ER, Pritchard JR, Church GM, Rubin EJ, Sassetti CM, Schnappinger D, Fortune SM. 2017. Programmable transcriptional repression in mycobacteria using an orthogonal CRISPR interference platform. Nature Microbiology 2:16274.

53. Rousset F, Cui L, Siouve E, Becavin C, Depardieu F, Bikard D. 2018. Genome-wide CRISPR-dCas9 screens in E. coli identify essential genes and phage host factors. PLoS Genet 14:e1007749.

54. Baba T, Ara T, Hasegawa M, Takai Y, Okumura Y, Baba M, Datsenko KA, Tomita M, Wanner BL, Mori H. 2006. Construction of Escherichia coli K-12 in-frame, single-gene knockout mutants: the Keio collection. Mol Syst Biol 2:2006.0008.

55. Yamamoto N, Nakahigashi K, Nakamichi T, Yoshino M, Takai Y, Touda Y, Furubayashi A, Kinjyo S, Dose H, Hasegawa M, Datsenko KA, Nakayashiki T, Tomita M, Wanner BL, Mori H. 2009. Update on the Keio collection of Escherichia coli single-gene deletion mutants. Mol Syst Biol 5:335.

56. Koo B-M, Kritikos G, Farelli JD, Todor H, Tong K, Kimsey H, Wapinski I, Galardini M, Cabal A, Peters JM, Hachmann A-B, Rudner DZ, Allen KN, Typas A, Gross CA. 2017. Construction and Analysis of Two Genome-Scale Deletion Libraries for Bacillus subtilis. Cell Syst 4:291–305.e7.

57. Sprenger GA. 1996. Carbohydrate metabolism in Zymomonas mobilis: a catabolic highway with some scenic routes. FEMS Microbiology Letters 145:301–307.

58. Geng B, Liu S, Chen Y, Wu Y, Wang Y, Zhou X, Li H, Li M, Yang S. 2022. A plasmid-free Zymomonas mobilis mutant strain reducing reactive oxygen species for efficient bioethanol production using industrial effluent of xylose mother liquor. Front Bioeng Biotechnol 10:1110513.

59. Torrents E. 2014. Ribonucleotide reductases: essential enzymes for bacterial life. Front Cell Infect Microbiol 4:52.

60. Kaiser M, Sawers G. 1994. Pyruvate formate-lyase is not essential for nitrate respiration by Escherichia coli. FEMS Microbiol Lett 117:163–168.

61. Knappe J, Wagner AF. 2001. Stable glycyl radical from pyruvate formate-lyase and ribonucleotide reductase (III). Adv Protein Chem 58:277–315.

62. Sutera VA, Han ES, Rajman LA, Lovett ST. 1999. Mutational analysis of the RecJ exonuclease of Escherichia coli: identification of phosphoesterase motifs. J Bacteriol 181:6098–6102.

63. Price MN, Wetmore KM, Waters RJ, Callaghan M, Ray J, Liu H, Kuehl JV, Melnyk RA, Lamson JS, Suh Y, Carlson HK, Esquivel Z, Sadeeshkumar H, Chakraborty R, Zane GM, Rubin BE, Wall JD, Visel A, Bristow J, Blow MJ, Arkin AP, Deutschbauer AM. 2018. Mutant phenotypes for thousands of bacterial genes of unknown function. 7706. Nature 557:503–509.

64. Cao Z, Mueller CW, Julin DA. 2010. Analysis of the recJ gene and protein from Deinococcus radiodurans. DNA Repair (Amst) 9:66–75.

65. Slade D, Radman M. 2011. Oxidative stress resistance in Deinococcus radiodurans. Microbiol Mol Biol Rev 75:133–191.

66. Deutschbauer A, Price MN, Wetmore KM, Tarjan DR, Xu Z, Shao W, Leon D, Arkin AP, Skerker JM. 2014. Towards an informative mutant phenotype for every bacterial gene. J Bacteriol 196:3643–3655.

67. Biegel E, Schmidt S, González JM, Müller V. 2011. Biochemistry, evolution and physiological function of the Rnf complex, a novel ion-motive electron transport complex in prokaryotes. Cell Mol Life Sci 68:613–634.

68. Mancini L, Terradot G, Tian T, Pu Y, Li Y, Lo C-J, Bai F, Pilizota T. 2020. A General Workflow for Characterization of Nernstian Dyes and Their Effects on Bacterial Physiology. Biophysical Journal 118:4–14.

69. Pirbadian S, Chavez MS, El-Naggar MY. 2020. Spatiotemporal mapping of bacterial membrane potential responses to extracellular electron transfer. Proc Natl Acad Sci U S A 117:20171–20179.

70. Jones CW, Doelle HW. 1991. Kinetic control of ethanol production by Zymomonas mobilis. Appl Microbiol Biotechnol 35:4–9.

71. Kalnenieks U. 2006. Physiology of Zymomonas mobilis: some unanswered questions. Adv Microb Physiol 51:73–117.

72. Rutkis R, Galinina N, Strazdina I, Kalnenieks U. 2014. The inefficient aerobic energetics of Zymomonas mobilis: identifying the bottleneck. J Basic Microbiol 54:1090–1097.

73. Yang S, Tschaplinski TJ, Engle NL, Carroll SL, Martin SL, Davison BH, Palumbo AV, Rodriguez M, Brown SD. 2009. Transcriptomic and metabolomic profiling of Zymomonas mobilis during aerobic and anaerobic fermentations. BMC Genomics 10:34.

74. Peters JM, Koo B-M, Patino R, Heussler GE, Hearne CC, Qu J, Inclan YF, Hawkins JS, Lu CHS, Silvis MR, Harden MM, Osadnik H, Peters JE, Engel JN, Dutton RJ, Grossman AD, Gross CA, Rosenberg OS. 2019. Enabling genetic analysis of diverse bacteria with Mobile-CRISPRi. Nature Microbiology 4:244–250.

75. Kuhns M, Trifunović D, Huber H, Müller V. 2020. The Rnf complex is a Na + coupled respiratory enzyme in a fermenting bacterium, Thermotoga maritima. 1. Communications Biology 3:1–10.

76. Schmehl M, Jahn A, Meyer zu Vilsendorf A, Hennecke S, Masepohl B, Schuppler M, Marxer M, Oelze J, Klipp W. 1993. Identification of a new class of nitrogen fixation genes in Rhodobacter capsulatus: a putative membrane complex involved in electron transport to nitrogenase. Mol Gen Genet 241:602–615.

77. Desnoues N, Lin M, Guo X, Ma L, Carreño-Lopez R, Elmerich C. 2003. Nitrogen fixation genetics and regulation in a Pseudomonas stutzeri strain associated with rice. Microbiology (Reading) 149:2251–2262.

78. Kremer TA, LaSarre B, Posto AL, McKinlay JB. 2015. N2 gas is an effective fertilizer for bioethanol production by Zymomonas mobilis. Proc Natl Acad Sci U S A 112:2222–2226.

79. Martien JI, Trujillo EA, Jacobson TB, Tatli M, Hebert AS, Stevenson DM, Coon JJ, Amador-Noguez D. 2021. Metabolic Remodeling during Nitrogen Fixation in Zymomonas mobilis. mSystems 6:e0098721.

80. Tatli M, Hebert AS, Coon JJ, Amador-Noguez D. 2019. Genome Wide Phosphoproteome Analysis of Zymomonas mobilis Under Anaerobic, Aerobic, and N2-Fixing Conditions. Front Microbiol 10:1986.

81. Khana DB, Tatli M, Rivera Vazquez J, Weraduwage SM, Stern N, Hebert AS, Angelica Trujillo E, Stevenson DM, Coon JJ, Sharky TD, Amador-Noguez D. 2023. Systematic Analysis of Metabolic Bottlenecks in the Methylerythritol 4-Phosphate (MEP) Pathway of Zymomonas mobilis. mSystems 8:e0009223.

82. Rudolf JD, Alsup TA, Xu B, Li Z. 2021. Bacterial terpenome. Nat Prod Rep 38:905–980.

83. Flesch G, Rohmer M. 1987. Growth inhibition of hopanoid synthesizing bacteria by squalene cyclase inhibitors. Arch Microbiol 147:100–104.

84. Horbach S, Neuss B, Sahm H. 1991. Effect of azasqualene on hopanoid biosynthesis and ethanol tolerance of Zymomonas mobilis. FEMS Microbiology Letters 79:347–350.

85. Li M, Hou F, Wu T, Jiang X, Li F, Liu H, Xian M, Zhang H. 2020. Recent advances of metabolic engineering strategies in natural isoprenoid production using cell factories. Nat Prod Rep 37:80–99.

86. Zhou J, Yang L, Wang C, Choi E-S, Kim S-W. 2017. Enhanced performance of the methylerythritol phosphate pathway by manipulation of redox reactions relevant to IspC, IspG, and IspH. J Biotechnol 248:1–8.

87. Kalnenieks U. 2006. Physiology of Zymomonas mobilis: some unanswered questions. Adv Microb Physiol 51:73–117.

88. Emms DM, Kelly S. 2019. OrthoFinder: phylogenetic orthology inference for comparative genomics. Genome Biol 20:238.

89. Emms DM, Kelly S. 2015. OrthoFinder: solving fundamental biases in whole genome comparisons dramatically improves orthogroup inference accuracy. Genome Biol 16:157.

90. Martin M. 2011. Cutadapt removes adapter sequences from high-throughput sequencing reads. 1. EMBnet.journal 17:10–12.

91. Bolger AM, Lohse M, Usadel B. 2014. Trimmomatic: a flexible trimmer for Illumina sequence data. Bioinformatics 30:2114–2120.

92. McClure R, Balasubramanian D, Sun Y, Bobrovskyy M, Sumby P, Genco CA, Vanderpool CK, Tjaden B. 2013. Computational analysis of bacterial RNA-Seq data. Nucleic Acids Res 41:e140.

93. Tjaden B. 2015. De novo assembly of bacterial transcriptomes from RNA-seq data. Genome Biol 16:1.

94. Tjaden B. 2020. A computational system for identifying operons based on RNA-seq data. Methods 176:62–70.

95. Virtanen P, Gommers R, Oliphant TE, Haberland M, Reddy T, Cournapeau D, Burovski E, Peterson P, Weckesser W, Bright J, van der Walt SJ, Brett M, Wilson J, Millman KJ, Mayorov N, Nelson ARJ, Jones E, Kern R, Larson E, Carey CJ, Polat İ, Feng Y, Moore EW, VanderPlas J, Laxalde D, Perktold J, Cimrman R, Henriksen I, Quintero EA, Harris CR, Archibald AM, Ribeiro AH, Pedregosa F, van Mulbregt P, SciPy 1.0 Contributors. 2020. SciPy 1.0: fundamental algorithms for scientific computing in Python. Nat Methods 17:261–272.

96. Bindels DS, Haarbosch L, van Weeren L, Postma M, Wiese KE, Mastop M, Aumonier S, Gotthard G, Royant A, Hink MA, Gadella TWJ. 2017. mScarlet: a bright monomeric red fluorescent protein for cellular imaging. Nat Methods 14:53–56.

